# Morphogen gradients are regulated by porous media characteristics of the developing tissue

**DOI:** 10.1101/2024.04.05.588250

**Authors:** Justina Stark, Rohit Krishnan Harish, Ivo F. Sbalzarini, Michael Brand

**Affiliations:** Technische Universität Dresden, Faculty of Computer Science, Germany; Max Planck Institute of Molecular Cell Biology and Genetics, Dresden, Germany; Center for Systems Biology Dresden, Dresden, Germany; Center for Regenerative Therapies Dresden, Dresden, Germany; Technische Universität Dresden, Center for Molecular and Cellular Bioengineering, Germany; Cluster of Excellence Physics of Life, TU Dresden, Germany

**Keywords:** morphogen gradient, extracellular space, complex geometry, image-based model, reaction-diffusion, gradient robustness, zebrafish, epiboly, fibroblast growth factor, GPU-accelerated simulation

## Abstract

Long-range morphogen gradients have been proposed to form by morphogen diffusion from a localized source to distributed sinks in the target tissue. The role of the complex tissue geometry in this process is, however, less well understood and has not been explicitly resolved in existing models. Here, we numerically reconstruct pore-scale 3D geometries of zebrafish epiboly from light-sheet microscopy images. In these high-resolution 3D geometries, we simulate Fgf8a gradient formation in the tortuous extracellular space. Our simulations show that when realistic embryo geometries are considered, a source-diffusion-degradation mechanism with additional binding to extracellular matrix polymers is sufficient to explain self-organized emergence and robust maintenance of Fgf8a gradients. The predicted normalized gradient is robust against changes in source and sink rates but sensitive to changes in the pore connectivity of the extracellular space, with lower connectivity leading to steeper and shorter gradients. This demonstrates the importance of considering realistic geometries when studying morphogen gradients.

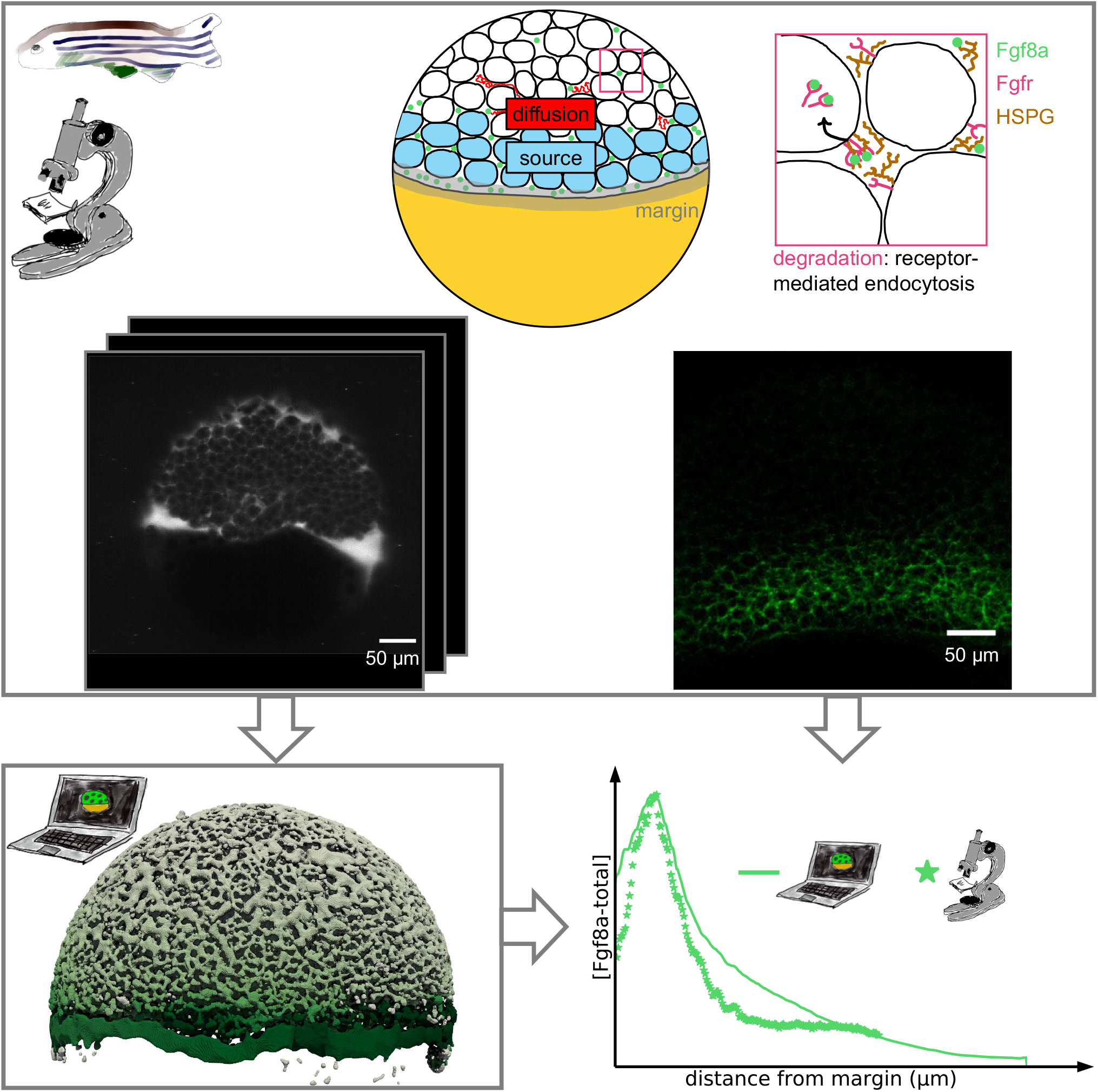

## 1 Introduction

The global body plan governing cell differentiation and tissue morphogenesis during embryonic development is defined by graded concentration fields of morphogens [33, 47, 99, 129]. Morphogens are signaling molecules, mostly proteins, secreted by localized source cells and moving through the embryo toward their target tissue, where they are degraded. Different modes of morphogen transport have been proposed, including diffusion, transcytosis, and directed transport via cytonemes [63, 83, 116, 125].

The simplest and most common mode of morphogen transport is by diffusion through the embryonic extracellular space (ECS) [26, 50, 133, 136]. The ECS is the complex-shaped interstitial space between the cells, which resembles a porous medium [86, 87]. When circumventing obstacles in tortuous porous media, molecules have to move further on average to reach the same mean square displacement as in free space, leading to reduced effective diffusivity [10, 14, 18, 20, 30, 39, 83, 106, 112]. Moreover, some parts of a porous medium may be inaccessible due to imperfect pore connectivity, i.e., dead-ends [53, 135], compartmentalizing the space.

Before diffusion through the ECS, morphogens are secreted by cells. Finally, cells take up extracellular morphogens by endocytosis. Both morphogen secretion and uptake amount to surface reactions, which depend on the surface area and accessibility of the cell surfaces. Pore accessibility and volume further influence bulk reactions such as morphogen interaction with the extracellular matrix [121]. Together, this results in a complex interplay of surface and bulk reactions, volume diffusion, and ECS geometry, as illustrated in Fig. 5a.

The porous ECS geometry can, therefore, be expected to significantly influence extracellular morphogen transport [106, 135]. The effective morphogen-receptor binding rate can be significantly smaller than the intrinsic rate and depend on the effective diffusivity [96, 126]. Diffusive hindrance, domain compartmentalization, and receptor localization hence influence morphogen gradient formation and maintenance in porous ECS geometries [70, 83, 120, 134, 135].

Another important factor modulating morphogen gradient formation is the transient binding of morphogens to non-receptors in the ECS, such as heparan sulfate proteoglycans (HSPGs) [4, 42, 69, 103, 116, 130]. HSPGs are tethered to cell membranes by a transmembrane core protein and have long poly-disaccharides reaching into the ECS, with heparan sulfate (HS) chains to which morphogens bind. HSPG binding has several effects on the morphogen gradient: in the ECS, it slows down morphogen diffusion while protecting morphogens from degradation by proteases [101]. At the cell membrane, it mediates complex formation with the morphogen receptor, locally enriching morphogens near cell surfaces. Concentrating morphogen diffusion to the cell surfaces and preventing morphogen loss and degradation in the ECS, HSPG binding stabilizes long-range gradients [49, 103, 130, 133]. All of these interactions add to the geometry influence and are subsumed in the *porous media characteristics* of the ECS.

A morphogen that forms long-range concentration gradients is Fibroblast growth factor 8a (Fgf8a). As a member of the highly conserved Fgf family, Fgf8a is involved in maintaining the midbrain-hindbrain boundary, somitogenesis, limb induction, and tissue patterning in zebrafish [32, 81, 97, 110]. In particular, Fgf8a forms a gradient along the animal-vegetal (AV) axis during zebrafish epiboly [50, 133].

Epiboly is the first morphogenetic movement in zebrafish during embryogenesis. It starts *≈*4 hours post fertilization (hpf) when actomyosin contraction in the yolk syncytial layer (YSL) pulls the enveloping layer (EVL) towards the vegetal pole [6, 22, 82], as depicted in Fig. 5b. This causes the blastoderm, consisting of the EVL and a deep-cell layer (DCL), to thin and spread over the yolk until completely engulfing the yolk by the end of epiboly [6, 22, 82].

During this dynamic cell movement, Fgf8a is secreted by a band of cells at the blastoderm margin, from where it diffuses throughout the ECS [50, 109, 133]. Interaction with HSPG at the cell surfaces and in the ECS is assumed to stabilize Fgf8a gradients by protecting Fgf8a from degradation and by slowing down its diffusion [49, 50, 133]. Cell-surface HSPGs also enhance Fgf8a binding to its receptors, Fgfr1 and Fgfr4, in the target tissue [49]. It is assumed that the internalized complex consists of Fgf8a, Fgfr, and HS in a 2:2:2 stoichiometry [88, 108]. Complex internalization involves clathrin-mediated endocytosis. Together, these processes—secretion, diffusion, degradation, and endocytosis—amount to a source-diffusion-degradation (SDD) [48, 63, 64, 116] mechanism with additional HSPG binding [50, 133].

Whether this SDD mechanism is sufficient to explain the experimentally observed Fgf8a gradient, however, has not been shown so far. It is difficult, if not impossible, to measure *in vivo* the individual roles of source and sink rates, HSPG binding, diffusion coefficient, and ECS geometry. To disentangle these different factors and quantify how they modulate morphogen gradient formation individually and collectively, mathematical and numerical models are required.

Many mathematical models of morphogen gradient formation have been developed over the decades [5, 8, 21, 27, 31, 43, 70, 75, 79, 84, 96, 124, 133]. Current models have, however, not explicitly resolved and studied the 3D porous geometry of the real embryo ECS. Instead, the diffusion space has been represented by simpler geometries, including 2D disks or 1D lines. 1D models have been successfully used to study morphogen gradient robustness to different source sizes [28] and to the absence of a sink [24], and how robustness can be established by non-receptor binding [69] and transcytosis [16]. 1D models have been further used to study scaling across different embryo sizes [43, 127], and local accumulation times (LATs) for different source [9] and sink configurations [21], as well as tissue growth [131].

Extensions of the LAT expressions to radially symmetric 2D and 3D models have shown that relaxation kinetics differ depending on the model dimensionality [46]. 2D models have further been used to study Dpp gradient precision [17] and the role of endocytic trafficking in Dpp gradient formation [16, 67], later extended to effective nonlinear transport equations with analytical expressions for concentration-dependent diffusion coefficient and degradation rate [15]. Models in symmetric and parametrizable geometries are popular for studying the influence of a single factor at a time and for deriving analytical solutions of morphogen gradients.

Another way to simplify the system for deriving analytical solutions is through geometrical upscaling, i.e., collectively representing the tortuous diffusion paths by an effective average constant [60]. Numerical homogenization is a common upscaling technique, which has been used to model morphogen gradient formation in the cortical region of the embryonic *Drosophila* syncytium [102], enabling the derivation of an analytical mean-field expression for that system.

Numerical solutions of models allow going beyond simple geometries and enable incorporating experimental image data. Image-based models of embryonic development have been developed in 2D+time, e.g., to model limb growth and digit patterning during vertebrate development [13, 71, 77, 79, 85] and kidney branching morphogenesis [78], as well as in 3D+time to model mouse limb-bud growth [29].

These image-based geometries, however, consisted of a singly connected surface (or outline in the 2D models), differing considerably from the porous zebrafish ECS geometry with its many intricately shaped and disconnected surfaces. These ECS surfaces, mostly cell surfaces, play an important role in gradient formation by confining the diffusion space and localizing morphogen sources and sinks. Despite this significant role, the zebrafish ECS geometry has been simplified in models of morphogen gradient formation during epiboly to the arc of a circle [83, 133] or the surface of a sphere [72, 73].

Simplified models are often well suited to test general principles of gradient formation. To better understand the influence of locally varying dynamics, anisotropies, and zonation effects, however, requires accounting for geometric heterogeneities. In addition, models of realistic embryo geometries are closer to the *in vivo* experiment, acquired in the real embryo geometry, against which they are validated.

The embryos’ spatial complexity and temporal deformation pose challenges in modeling their realistic geometries. As embryo geometries cannot be mathematically parameterized, they require algorithmic representations at high resolution to capture pore-scale dynamics. In particular, morphogen gradient formation is a molecular process influenced by the ECS geometry across several orders of magnitude, from interstitial spaces (nanometers) to the embryo size (millimeters). Computational models that consider all length scales can be prohibitively expensive to compute. Memory-efficient algorithms, like geometry-adaptive sparse grids [57], and acceleration of the multi-scale simulation in Graphics Processing Units (GPUs) are thus required to render simulations in realistic embryo geometries feasible.

Here, we use a memory-efficient GPU-accelerated algorithm [117] to address the above biological questions by simulating Fgf8a gradient formation in realistic 3D ECS geometries of zebrafish epiboly derived from light-sheet microscopy volumes at the resolution of individual cells. In these image-based geometries, we simulate morphogen gradient formation by solving a system of coupled partial differential equations, accounting for the spatial variations of sources and sinks and the interaction of Fgf8a with HSPG. Using this image-based model, we quantify gradient robustness to changes in different factors, such as source and sink rates, HSPG binding, morphogen diffusivity, and ECS geometry.

We find that the normalized Fgf8a gradient is robust against changes in the source and sink rates, whereas it is sensitive to changes that affect diffusive hindrance. Besides changes in the molecular diffusion constant of the morphogen, the latter includes changes in the extracellular HSPG concentration and changes to the ECS geometry. Concretely, peak concentrations near the source are more preserved, and the gradients are steeper and shorter for higher HSPG concentrations, lower diffusion coefficients, and higher ECS tortuosity.

## 2 Materials and Methods

We state the equations modeling Fgf8a gradient formation, discuss the numerical method and parallel computing software used to numerically solve these equations, and we describe image acquisition and image processing.

### 2.1 Model equations

Fgf8a gradient formation is explained by an SDD mechanism with HSPG binding [50, 133]. Specifically, two diffusing fractions of Fgf8a with different diffusion coefficients have been identified *in vivo*: a major fraction of *≈*93 % moving freely with *D* = 55 µm^2^s^−1^ and a HSPG-bound fraction of *≈*7 % moving with *D* = 4 µm^2^s^−1^ [50]. In addition to interacting with extracellular HSPG, Fgf8a binds to cell-surface HSPG, which mediates its receptor complex formation [49, 88, 108]. The cell-surface bound Fgf8a fraction is not detected by single-molecule fluorescence correlation spectroscopy (FCS) measurements in the ECS but by Heparinase (HepI) injection experiments, in which cleavage of the HS side chains has been shown to lead to an increase in overall Fgf8a concentrations detected by FCS [50].

Based on these experimental findings, we account for the different Fgf8a interactions by distinguishing four concentration fields in space and time: freely diffusing Fgf8a ([Fgf8a]), slowly diffusing HSPG-bound Fgf8a in the ECS ([Fgf8a : HSPG^ECS^]), immobile cell-surface HSPG-bound Fgf8a ([Fgf8a : HSPG^cellSurf^ ]), and the immobile ternary 2:2:2 complex of Fgf8a, Fgfr, and HSPG at the cell surface

([Fgf8a_2_ : 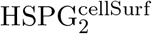: Fgfr_2_]). The biochemical kinetics between these species in the ECS is:

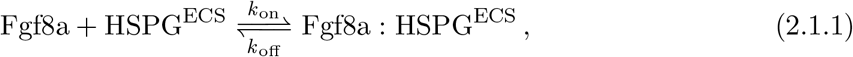

at the cell surfaces, it is:

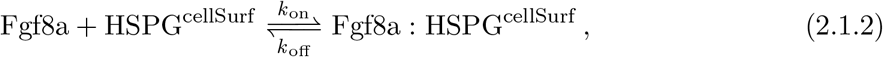

and ternary receptor complex-formation occurs as:

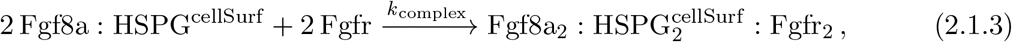

where *k*_on_ and *k*_off_ are the association and dissociation rates of Fgf8a to HSPG, and *k*_complex_ is the association rate of the ternary [Fgf8a_2_ : 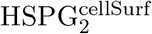: Fgfr_2_] complex. As cell-surface HSPGs have been shown to trap Fgf8a and mediate receptor complex-formation and dimerization [49, 88], our model assumes that Fgf8a binding to HSPG^cellSurf^ precedes complex formation. Further, the model combines complex association, dissociation, and dimerization into a single effective forward reaction to reduce the number of unknown rate constants.

Applying the principle of mass-action and assuming diffusive transport, these kinetics yield the following partial differential equations for the evolution of the four concentration fields in space ***x*** and time *t*:

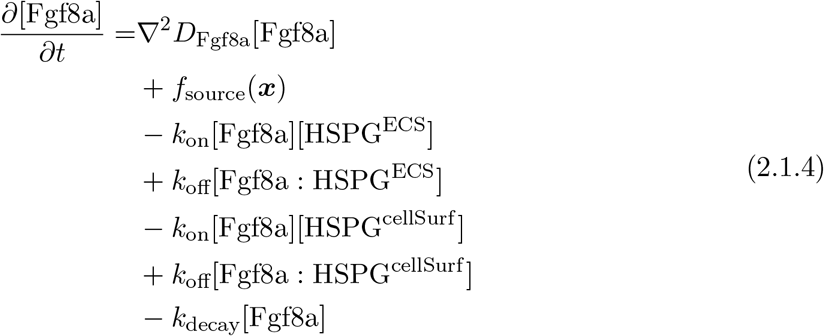

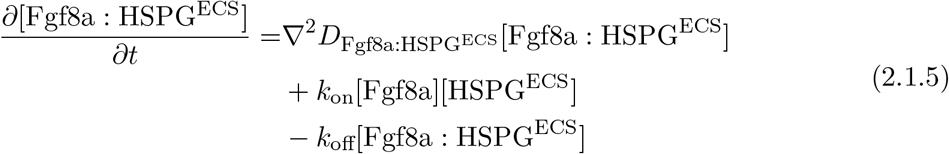

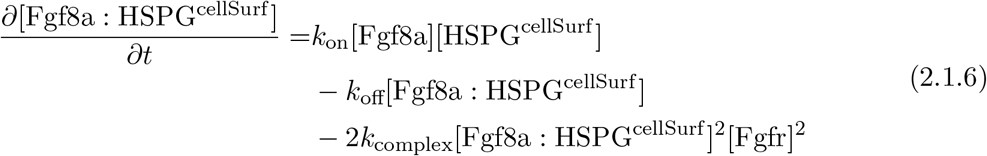

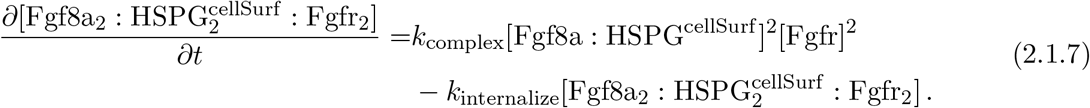

*D*_Fgf8a_ and *D*_Fgf8a_:HSPG^ECS^ are the diffusion coefficients of the free and HSPG-bound Fgfa8 fractions in the ECS, respectively. Both diffusion coefficients are constant in space and time, neglecting spatial variations of extracellular matrix viscosity. [HSPG^ECS^] and [HSPG^cellSurf^ ] are the HSPG concentrations with free binding sites in the ECS and at the cell surfaces, respectively. The rate *k*_decay_ describes the decay of free Fgf8a in the ECS through thermal denaturation and proteolysis [65, 101]. The rate *k*_internalize_ is the complex internalization rate through endocytosis. The model assumes that for each internalized complex, two new Fgfr and HSPG^cellSurf^ appear at the cell membrane, respectively, i.e., that their total concentration is constant over time. This simplifies the more complicated real receptor fate, of which Fgfr4 has been shown to be mainly recycled, whereas Fgfr1 with ligand is mainly degraded in the lysosome [51]. Fgf8a secretion by source cells is represented by the source term *f*_source_(***x***).

Fgf8a sources and sinks vary in space. Concretely, only the deep cells participate in Fgf8a production and uptake, i.e., there is no production and uptake in the YSL and EVL (see Fig. 5b). We denote ECS boundaries that belong to either the EVL or YSL by ∂Ω_shell_, such that the surfaces of the deep cells are defined by ∂Ω_ECS_*\*∂Ω_shell_. Within the deep cells, we further spatially restrict the source to a band of cells near the blastoderm margin. We define the width of this cell band *w*_source_ and its distance from the margin *d*_margin_, so that we can mathematically formalize the source term as

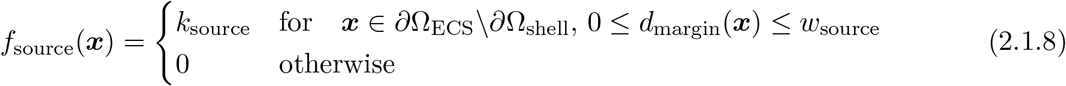

with source rate *k*_source_.

In addition to Fgf8a molecular degradation with rate *k*_decay_ everywhere in the ECS, ternary complex formation and subsequent endocytic uptake constitutes an additional Fgf8a sink *f*_sink_(***x***). This endocytic sink is localized only to the deep cell surfaces ∂Ω_ECS_*\*∂Ω_shell_ by setting [Fgfr] to zero everywhere else. We model *f*_sink_(***x***) as a first-order reaction with rate *k*_internalize_

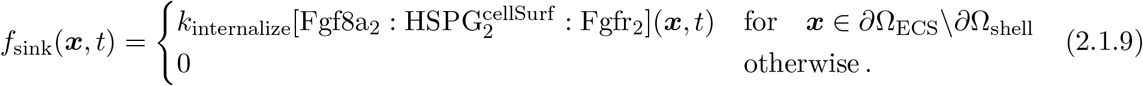

### 2.2 Numerical method

To numerically model localized sources and sinks and solve Eq. (2.1.4)–(2.1.7) in pore-scale 3D ECS geometries derived from images, we use an open-source image-based simulation pipeline for reaction-diffusion processes in porous media [117]. This pipeline computes a level-set representation of the diffusion domain based on volumetric microscopy images or microcomputed tomography scans. Based on the level-set function, it generates geometry-adapted sparse grids on which it solves inhomogeneous reaction-diffusion partial differential equations with multi-GPU acceleration. A detailed explanation of the numerical method has been given by Stark and Sbalzarini (2023).

The pipeline addresses the high memory requirements of multi-scale models using geometry-adapted sparse grids [57], enabling individual point allocation in any domain subspace as needed. As sparse grids preserve the Cartesian neighborhood of a structured grid, they enable fast neighborhood access and differential operator approximation using standard grid-based schemes. Sparse grids are, therefore, suitable for discretizing porous media geometries in a memory- and compute-efficient way.

Generating such a geometry-adapted sparse grid for the complex-shaped geometry of an embryonic ECS diffusion domain Ω requires a numerical description of its boundary ∂Ω embedded in the grid, i.e., the interface between Ω and the surrounding space *S*. We use the level-set method [89, 118] to implicitly represent ∂Ω as the zero-level set of a higher-dimensional function *ϕ*. Specifically, we use the signed-distance function (SDF):

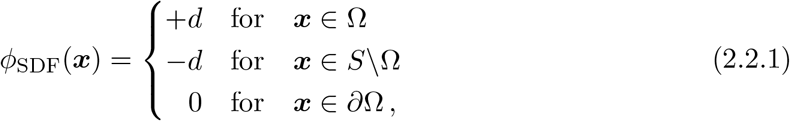

which is smooth around the surface ∂Ω and provides geometric information through the orthogonal distance *d* of any point ***x*** in the domain to the closest point on the nearest surface.

We compute *ϕ*_SDF_ by first generating a binary pixel mask of the ECS from the image using the pixel-classification tool ilastik [11]. This represents is the indicator function for Ω. Using this indicator function, we initialize *ϕ*(***x***, 0) with +1 inside Ω and -1 outside Ω. This serves as initial condition for computing *ϕ*_SDF_ using the classic Sussman level-set redistancing algorithm [118], which finds the SDF as the stationary solution of

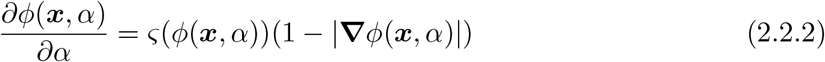

for large *α*. To prevent interface shifting across cell boundaries for steep *ϕ*, and to accelerate convergence of the iterations if *ϕ* is flat near the interface, the smoothed sign function *ς*is chosen as [91]:

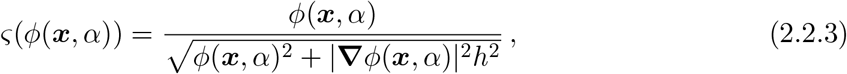

where *h* is the uniform grid spacing.

We perform Sussman redistancing with multi-CPU parallelization on a dense grid with first-order accuracy in space and time [117]. Using the so-computed *ϕ*_SDF_, a geometry-adapted sparse grid containing only points in the diffusion domain is generated to reduce memory requirements. This sparse grid stores the four concentration fields along with *ϕ*_SDF_. Together, this enables defining spatially varying diffusion coefficients and reaction rates, including surface reactions. Using *ϕ*_SDF_, which indicates on which side of the surface a grid point is located, we impose zero-flux Neumann boundary conditions within the finite-difference stencil.

Taking advantage of the regular structure of sparse grids, the diffusion operators are discretized using finite-difference methods, which afford efficient memory access and parallelize well on multi-CPU and multi-GPU, in particular when an explicit time-stepping scheme is used [25, 61, 132]. Since we here consider a strongly memory-limited problem, we solve the model in time using the first-order explicit Euler time-stepping scheme, which does not require storing and communicating intermediate stages.

### 2.3 Software implementation

Even when using geometry-adapted sparse grids, the pore-scale models of the embryonic ECS exceed single-processor capacity by far, requiring efficient parallelization. We use the open-source C++ library OpenFPM for parallelizing the simulations [59]. OpenFPM data structures, which are generated at compile time using template meta-programming, hide communication from the application and are portable across CPU and GPU architectures [58].

OpenFPM implements distributed sparse block grids by dividing the domain into chunks of 8 *×* 8 *×* 8 grid points and only allocating chunks containing at least one grid point [57]. This considerably reduces memory requirements for sparse geometries, such as the zebrafish ECS. OpenFPM sparse grids are also multi-GPU accelerated, significantly reducing simulation times compared to simulations on multi-CPU [57]. These simulations have been shown to scale with problem size and number of GPUs for irregular porous media geometries [117]. In particular, it has been shown that the sparser the geometry and the locally denser the grid points, the larger the savings in memory and simulation run time [57, 117]. The zebrafish ECS has a particularly sparse geometry, occupying only *≈*4% of 3D light-sheet microscopy volume, making it well-suited for OpenFPM sparse-grids discretization.

We visualized simulation results using Paraview [2]. Plots were generated using the Matplotlib library [55].

### 2.4 Acquisition of light-sheet microscopy time-lapse video

To geometrically characterize the zebrafish ECS and its boundaries, we collected Tg(bactin:hRas-EGFP) embryos, in which Enhanced Green Fluorescent Protein (EGFP) was targeted to the cell membrane. As the embryos reached the sphere stage, they were dechorionated and injected with 0.2 nL of a 5 µM solution of Tetramethylrhodamine (TMR)-labeled dextran at their animal pole, where ECS is abundant. As a hydrophilic polysaccharide, dextran does not cross the membrane barrier and distributes throughout the ECS as the embryo develops. This enabled us to track the ECS over the course of epiboly.

To perform multi-view imaging of embryos for 3D modeling, we used the Plan-Apochromat 20X/1.0W objective of the Zeiss light-sheet Z.1 microscope. For this, we first transferred the TMR-Dextran-injected embryos into a tube containing 1% low-melting point agarose, mixed with F-Z fluorescent microspheres at a 1:4000 dilution, stored at 42°C to prevent solidification. Together with the agarose–bead mix, the embryos (maximum three) were then sucked into a glass capillary (20 µL volume, 1 mm diameter) by pulling on the inserted plunger, such that the embryos were all spaced evenly inside the capillary. Once the agarose solidified, we transferred the capillary to a beaker containing E3 medium until the imaging chamber at the microscope was assembled. Thereafter, the capillary was inserted into the imaging chamber (also filled with E3), positioned, and automatically detected by imaging. An embryo embedded in the solidified agarose–bead mixture was then slowly pushed out of the capillary using the plunger until it reached the center of the detection volume.

Light-sheet adjustment, multi-view image-acquisition, and later multi-view reconstruction were performed according to the protocol detailed by Icha et al. (2016) using the Fiji Big-DataViewer plugin [94, 95]. Image acquisition was performed from five different angles over the course of epiboly. The resulting time-lapse multi-view image data set is composed of 25 frames of 3D image volumes acquired from 5 hpf (*≈*40% epiboly) to 9 hpf (*≈*90% epiboly) (voxel size = 0.4574 µm in all three dimensions, stack size: 1834*×*1739*×*1551 pixel). This allowed visualizing changes in the spatial arrangement of cells and the geometry of the ECS over the course of early development. Fig. 5c shows exemplary z-slices of the video with the EGFP-hRas-labeled cell membranes in green and the TMR-Dextran-labeled ECS in magenta.

### 2.5 Reconstructing 3D zebrafish ECS geometries based on light-sheet microscopy images

Using the TMR-dextran fluorescence signal of the light-sheet time-lapse video, we modeled ECS geometries using level-set sparse grids as explained in Sections 2.3 and 2.2. This image-based ECS reconstruction consisted of five steps, the intermediate results of which are shown in Fig. 1a– e for an exemplary z-slice. The steps were: 1) extracting volumes of individual time points from the light-sheet video; 2) image processing and segmentation using Fiji [107] and ilastik [11]; 3) processing of the segmentation result and conversion to an indicator function; 4) computing a numerical surface description as a signed distance function (SDF) using the level-set method; 5) generating a geometry-adapted discretization of the ECS using distributed sparse block grids [57]. We next describe these five image-based modeling steps in more detail.

**Figure 1.**
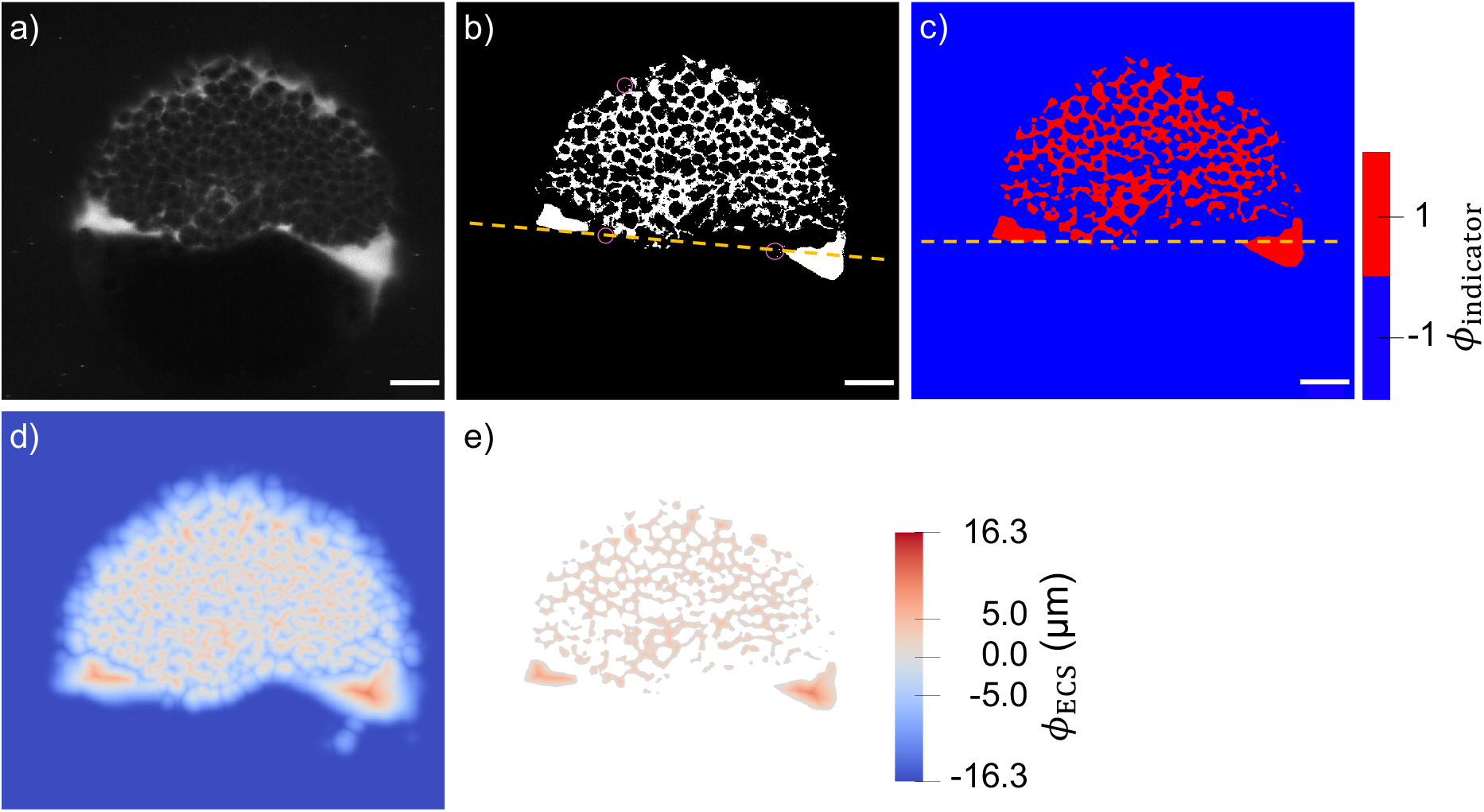
Steps for modeling ECS geometries from light-sheet microscopy volumes visualized for an exemplary time point (12 out of 25, *≈*60 % epiboly) and z-slice (plane 480 from the epiblast in the direction of hypoblast and the yolk, i.e., the mid-plane formed by the AV/DV axes). a) TMR-dextran channel of the light-sheet data after 3D multi-view fusion; fluorescence marks ECS. b) Segmentation mask of the ECS (white) with a yellow dashed line showing the margin plane and pink circles highlighting specks of noise and small unconnected islands that were subsequently removed. c) Binary indicator function of the ECS after mask denoising and horizontal alignment of the margin plane (yellow dashed line). d) Redistanced signed-distance function *ϕ*_ECS_ of the level-set representation of the ECS surface. e) Sparse block grid storing only points in the ECS; scale bars: 50 µm.

The light-sheet video (exemplary z-slices in Fig. 5c) was stored as a 0.7 TB HDF5 file containing five viewpoints plus their 3D fusion for all 25 time points. For geometry modeling the geometry, 3D fused views were extracted as separate files to reduce RAM/DRAM requirements. Extraction of the 3D multiview reconstruction of the TMR-dextran channel at each time point was done using the Multiview Reconstruction Tool from the Fiji BigDataViewer plugin [94, 95]. An exemplary z-slice is shown in Fig. 1a.

The resulting TIFF files were subsequently converted to the ilastik HDF5 file format using the ilastik Import/Export plugin in Fiji. In ilastik [11], the interactive Random Forest pixel classifier was used to obtain a pixel mask for the ECS. The classifier was trained by manual image example annotation for the two pixel classes, ECS versus non-ECS, where non-ECS included pixels of background, cells, and yolk. Pixel class annotation and prediction were repeated for different *x, y*, and *z*-slices of the 3D image volume at video time point 12 until the classification results no longer improved. The trained classifier was then used for pixel classification of the other 24 time points of the video without further manual annotation.

The segmentation result, shown in Fig. 1b for an exemplary z-slice, likely overestimates the ECS volume because ECS structures can be below pixel resolution. Nevertheless, the network character of the ECS geometry was successfully captured.

Postprocessing of the segmentation mask (pixel value of 2 in the ECS and 1 otherwise) was done before conversion to an indicator function. This included Gaussian filtering (*σ* = 2) and alignment of the sample rotation using the `scipy.ndimage` Python package [122]. Sample alignment made the margin plane horizontal (yellow dashed line in Fig. 1b and c) by rotation of the 3D pixel mask about the z-axis, applying the standard rotation matrix using trigonometric functions. The rotated mask was then interpolated back to the pixel grid using third-order spline interpolation. This was followed by thresholding the filtered and rotated segmentation mask, which was no longer binary, setting pixels to one with value *>* 1.4 and all others to −1. The threshold was determined such that it removes unconnected islands of size ≤ 3 *×* 3 *×* 3 pixels, whose maximum value after Gaussian smoothing was 1.39 (examples marked by pink circles

in Fig. 1b. This denoising step was necessary to avoid numerical instabilities in the Sussman algorithm that would otherwise be caused by imaging noise, at the expense of introducing a small bias and artificial smoothing of the ECS surface. This resulted in an indicator field as exemplary shown in Fig. 1c, which was stored as a binary file. Each of these binary files was 162.5 GB in size. The indicator function was then loaded onto a distributed-memory 3D computational grid on a compute cluster with multiple CPUs. Thin structures *<* 2 pixel in width were removed to fulfill the level-set resolution requirement resulting from the Nyquist-Shannon sampling theorem [117]. Finally, a representation of the ECS surfaces was computed as a SDF *ϕ*_ECS_, as explained in Section 2.2. The exemplary result is shown in Fig. 1d. Based on this *ϕ*_ECS_, a sparse block-grid representation of the ECS was generated, resulting in a memory-efficient geometry-adapted data structure as illustrated in Fig. 1e. A full 3D visualization of the resulting surface is shown in Fig. 5 d – g for the time point at *≈*60% epiboly. Following this procedure, we built models of the ECS geometries for all 25 time points of the light-sheet time-lapse video. Visualizations of the results at all time points are shown in Suppl. Fig. S9.

At full resolution, with one grid point per light-sheet microscopy voxel, representing the zebrafish ECS on a sparse block grid reduced memory requirements by a factor of 17, from *≈*306 GB (dense grid, no blocks) to *≈*17 GB (sparse block grid). This enables fully resolved numerical simulations on a GPU, since the sparse-grid data fits the DRAM of a typical GPU (here: Nvidia A100 with 40 GB DRAM). We further found that simulation results at full and half grid resolution were identical, indicating that the pixel resolution of the geometry was sufficient to yield grid-converged numerical results. Using half the resolution resulted in an overall 28-fold reduction in simulation times (from 110 hours to 4 hours of GPU compute time per run simulating 60 minutes of physical time). We therefore used a half-resolution computational grid (i.e., *h* = 2 pixel) in all simulations, resulting in a grid spacing of 0.92 µm and a memory requirement of 3.5 GB.

As the time resolution of the light-sheet microscopy video was not fine enough to allow for the derivation of continuous ECS deformation maps, we simulated gradient formation in fixed ECS geometries, neglecting advective transport and growth. We imposed zero-flux Neumann boundary conditions at all interfaces, assuming that the fluid inside the ECS does not permeate the cell membranes. With these assumptions and using the numerical methods from Section 2.2, we solved the model equations from Section 2.1 in the 3D ECS geometries at *≈*40, 60, and 75% epiboly to study morphogen gradient emergence and maintenance.

### 2.6 Modeling localized sources and sinks

In order to spatially restrict sources and sinks according to Eqs. (2.1.8) and (2.1.9) to the surfaces of the deep cells, these surfaces need to be distinguished from those of the YSL and EVL, i.e., what we call the *shell* boundary ∂Ω_shell_. To accomplish this, we computed another SDF, *ϕ*_shell_, representing ∂Ω_shell_, following the procedure as described in Section 2.5, but using segmentation masks in which the cells have been blurred out by Gaussian filtering (*σ* = 3) followed by setting pixels to one whose value *>* 1.0 and all others to −1.

This resulted in an SDF as shown in Fig. 2a. Combining *ϕ*_shell_ and *ϕ*_ECS_ enables defining the deep-cell surfaces as the set of grid points 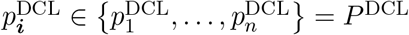 within one grid spacing from ∂Ω_ECS_ and further than 15 µm away from ∂Ω_shell_, i.e., *P*^DCL^:= {*i* ∈ℕ^3^: 0 *< ϕ*_ECS_(***x***_***i***_) *< h* and *ϕ*_shell_ to which we restrict [Fgfr] and [HSPG^cellSurf^ ].

**Figure 2.**
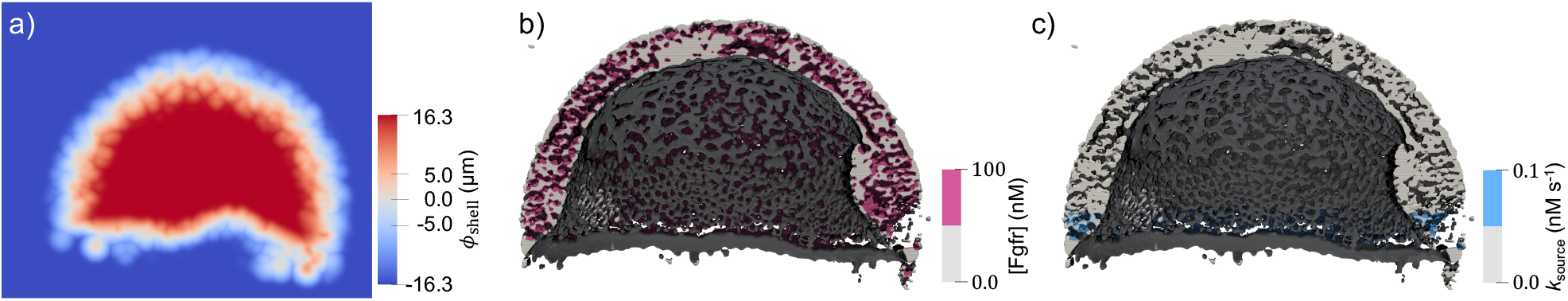
Spatially varying sources and sinks in the 3D model at an exemplary time point at *≈*60% epiboly. a) SDF of the embryo shell boundary, *ϕ*_shell_, for an exemplary z-slice. Combining *ϕ*_shell_ (representing YSL and EVL boundaries) with *ϕ*_ECS_ enables restricting sources and sinks to the deep-cell surfaces. Clipped model of the ECS showing that Fgf-receptors ((b) in pink) are restricted to the deep cell boundaries, i.e., not EVL nor YSL. The sources ((c) in blue) are restricted to *≈*5 rows of deep cells above the margin of the blastoderm. Concentrations are given in units of nM, *k*_source_ in 1/s.

Fgf8a secretion was also restricted to this same set of grid points, additionally restricting the source location to a band of deep cells above the blastoderm margin [109]. The width of this band was *≈*5 cells at early gastrula [50] and *≈*8 cells at mid gastrula [109]. For this, *w*_source_ in Eq. (2.1.8) was set to 70 µm at 40 and 60% epiboly and to 112 µm at 70% epiboly, assuming an average cell diameter of 14 µm [50]. The model assumes that source cells also act as sink cells. Comparing the simulation results of this model with a model that assumes source cells do not act as sinks, we found that the normalized gradient profiles are unaffected by this assumption (see Supplementary Note S3). The resulting sources and sinks of the model are visualized in Fig. 2 for an exemplary time point at *≈*60% epiboly.

### 2.7 Parameter search

A major difficulty of modeling morphogen gradient formation realistically, such that it can be validated with experimental data, is that various required key parameters remain unknown because they are difficult or impossible to measure experimentally. These include morphogen production and degradation rates, receptor and non-receptor binding affinities and concentrations, and advection velocities due to growth. The models hence remain underdetermined with multiple unknown parameters, requiring parameter searches and fitting [114].

The model described in Eqs. (2.1.4)–(2.1.7) has nine unknown parameters, in particular, [Fgfr], [HSPG^ECS^], [HSPG^cellSurf^ ], *k*_on_, *k*_off_, *k*_decay_, *k*_complex_, *k*_internalize_, and *f*_source_. We performed a systematic parameter search to find values of the parameters that maintain the *in vivo* gradient.

To maximize the predictive power of our model, we fixed as many parameters as possible to experimental or literature values, as summarized in Table 1. One of the unknown parameters is the overall binding and unbinding rate of Fgf8a to HSPG^ECS^ and HSPG^cellSurf^. For the two HSPG families, glypican and syndecan, the dissociation constants, defined as 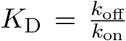, of the Fgf8-HSPG interaction 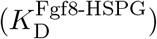 have been determined by *in vivo* dual-color fluorescence cross-correlation spectroscopy (FCCS) in the gastrulating zebrafish: *K*_D_ = 1.03 1.24 µM (syndecans) and *K*_D_ = 6.15 *±* 0.74 µM (glypicans) [49]. Hence, both affinities are more than an order of magnitude lower than Fgf8-receptor binding affinities (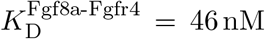 and 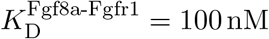 [98]).

**Table 1:** Summary of all model parameters. We combined literature values (references) with own experimental measurements (“Exp.”) and deductive reasoning from known values as described in the text (“Text”). The remaining three parameters were determined in a computational parameter search (“Search”).

|  |  |  |  |
| --- | --- | --- | --- |
| Diffusion coefficients |  |  |  |
| Fgf8a (93 % of extracellular Fgf8a) | $D_{\text{Fgf8a}}$ | $55 \mu\text{m}^2\text{s}^{-1}$ | [50] |
| Fgf8a:HSPG (7 % of extracellular Fgf8a) | $D_{\text{Fgf8a:HSPG}^{\text{ECS}}}$ | $4 \mu\text{m}^2\text{s}^{-1}$ | [50] |
| Concentrations |  |  |  |
| maximal concentration of extracellular Fgf8a |  | 7.9 nM | Exp. |
| extracellular HSPG concentration | $[\text{HSPG}^{\text{ECS}}]$ | 0.4 $\mu\text{M}$ | Search |
| cell-surface HSPG concentration | $[\text{HSPG}^{\text{cellSurf}}]$ | 1.0 $\mu\text{M}$ | Search |
| Fgf receptor concentration | $[\text{Fgfr}]$ | 0.1 $\mu\text{M}$ | Text |
| On and off binding rates |  |  |  |
| Association rate of Fgf8a and $\text{HSPG}^{\text{ECS}/\text{cellSurf}}$ | $k_{\text{on}}$ | $4.6 \times 10^{-6} (\text{nM s})^{-1}$ | Text |
| Dissociation rate of Fgf8a and $\text{HSPG}^{\text{ECS}/\text{cellSurf}}$ | $k_{\text{off}}$ | $0.023 \text{s}^{-1}$ | [41, 115] |
| Dissociation constant of Fgf8a : $\text{HSPG}^{\text{ECS}/\text{cellSurf}}$ | $K_{\text{D}}^{\text{Fgf8a:HSPG}}$ | 5.0 $\mu\text{M}$ | [49], Text |
| Association rate of Fgf8a : $\text{HSPG}^{\text{cellSurf}}$ and Fgfr | $k_{\text{complex}}$ | $4.6 \cdot 10^{-5} (\text{nM s})^{-1}$ | Text |
| Source and sink rates |  |  |  |
| Fgf8a production rate | $k_{\text{source}}$ | 0.1 $\text{nM s}^{-1}$ | Search |
| Fgf8a molecular decay rate | $k_{\text{decay}}$ | $1.3 \cdot 10^{-3} \text{s}^{-1}$ | [133] |
| Internalization rate for Fgf8a<sub>2</sub> : $\text{HSPG}_2^{\text{cellSurf}}$ : Fgfr<sub>2</sub> | $k_{\text{internalize}}$ | $9 \cdot 10^{-3} \text{s}^{-1}$ | Text |

Assuming that the syndecans and glypicans affinities are representative of the overall Fgf8a : HSPG interaction, and considering that more glypicans than syndecans are present in the zebrafish ECS [49], we set the dissociation constant of Fgf8a : HSPG^ECS^ and Fgf8a : HSPG^cellSurf^ in our model to an average 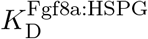 of 5.0 µM. Using this 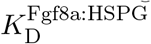 and an HSPG unbinding rate *k*_off_ = 0.023 s^−1^, as has been measured for Fgf2 [41, 115], we find the binding rate 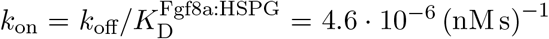. Further, we assume a 10-times higher Fgfr binding rate of *k*_complex_ = 4.6· 10^−5^ (nM s)^−1^ and set [Fgfr] = 0.1 µM by approximating volume concentrations from experimentally determined Fgfr surface concentrations [98].

It has been experimentally shown that Fgf8a concentrations are higher at the membrane than in the ECS [49] and that cleavage of HS side-chains upon heparinase injection causes a two-fold increase in Fgf8a concentration levels in the ECS [50]. The exact factor by which [HSPG^cellSurf^ ] is higher than [HSPG^ECS^] is, however, unknown, so we set it arbitrarily to 2*×*.

From the experimentally measured *k*_decay_ = 1.3· 10^−3^ s^−1^ [133], and taking into account geometrical hindrance as discussed in Supplementary Note S5 and the gradient half-decay length *λ* from the *in vivo* gradient profile in Fig. 7e, we estimate

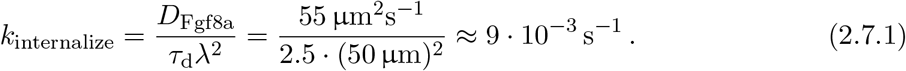

For the computational parameter search, we assumed that the experimentally measured intensity profile at 60% epiboly was at a steady state for this time point. We then searched for values of the remaining three parameters (“Search” in Table 1) that maintained this steady-state profile. For this, we initialized the Fgf8a concentration in the geometry reconstruction at 60% epiboly with the experimental profile shown in Fig. 3a as detailed in Supplementary Note S1. To compare the 3D simulation results with the 1D experimental gradient, we computed 1D profiles along the AV axis by averaging the sum of all four Fgf8a fractions in each *xz*-plane of the simulation grid. These concentration averages were subsequently normalized and plotted against the distance from the blastoderm margin, as shown in Fig. 3b. Using the experimental profile as initial condition, we computed the solution of Eqs. (2.1.4)–(2.1.7) for different parameter sets until *t*_max_ = 60 min, which was sufficiently long to ensure that gradients have reached a steady With these assumptions, and using the experimental profile as initial condition, we tested different *k*_source_ and HSPG concentrations in simulations. These two parameters mainly influenced the Fgf8a / Fgf8a : HSPG^ECS^ ratio in the simulations, i.e., the higher [HSPG], the higher the bound fraction. We found that [HSPG^ECS^] = 0.4 µM and [HSPG^cellSurf^ ] = 1.0 µM closely matched the experimentally measured Fgf8a / Fgf8a : HSPG^ECS^ ratio (simulated ratio: 92.6% / 7.4%) after 60 minutes of simulated time, see Fig. 4b. For these parameter values, the concentration levels of the overall Fgf8a profile were well maintained when using a source rate of 0.1 nM s^−1^, as shown in Fig. 4a.

**Figure 3.**
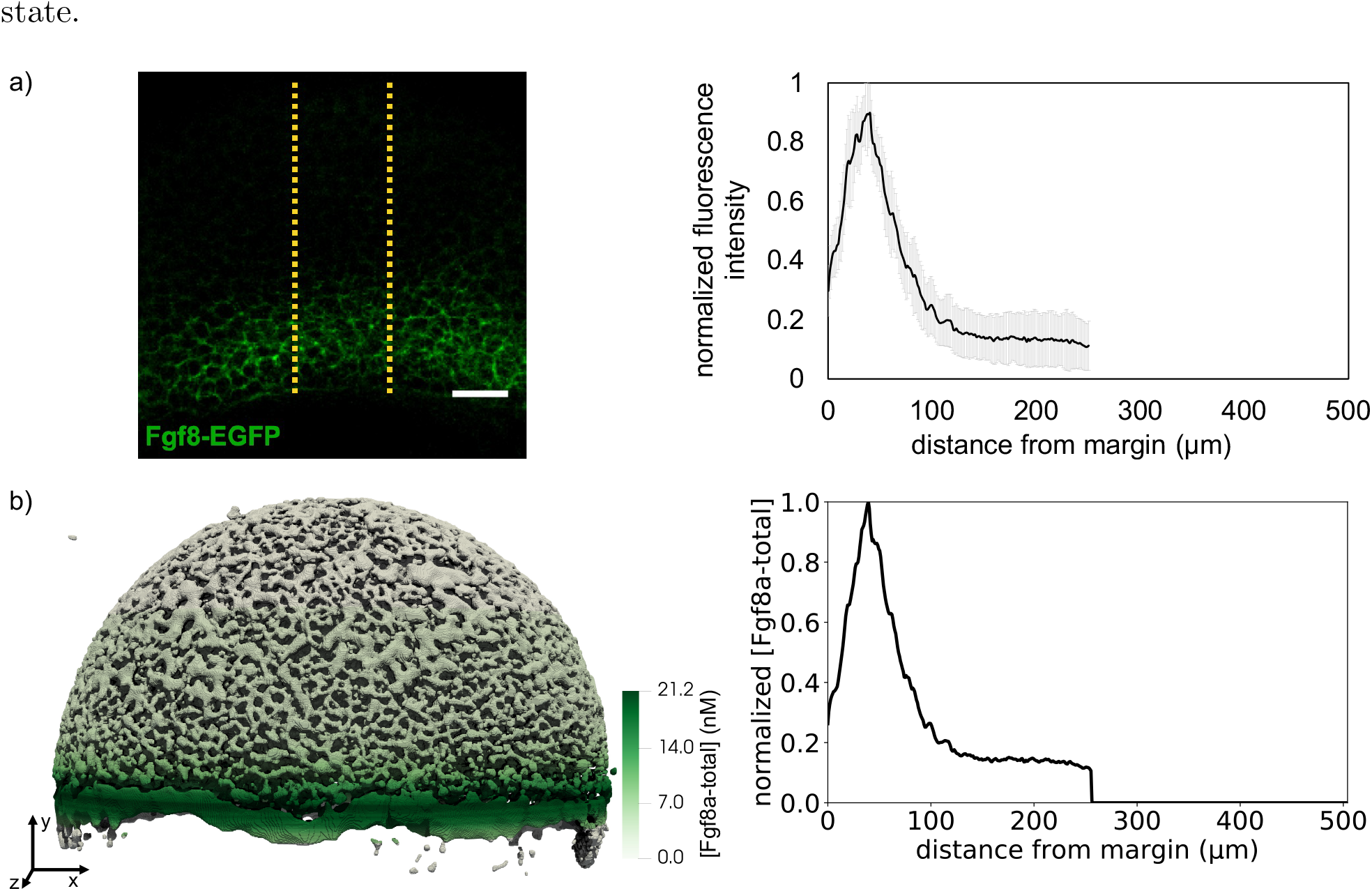
Initialization of the total Fgf8a concentration with the experimental fluorescence intensity profile from at *≈*60% epiboly. a) The fluorescence intensity profile (right) is extracted from sum-intensity z-projected confocal images (left). Values are intensity averages across a 100 µm wide region (neural plate). Figure reproduced with permission from Harish et al. (2023). The embryo is oriented as in Fig. 5b and depicted in the inset (ventral left, dorsal right, animal pole top, vegetal pole bottom). Scale bar: 50 µm. Data are represented as mean *±* standard deviation. b) The concentration field in the simulation (left) was obtained by linearly interpolating between the discrete margin distances of the experimental profile and uniformly distributing the mass in each *xz*-plane. We converted normalized intensity values to concentrations by scaling with the maximum concentration measured by FCS. For comparison with experiments, we computed 1D profiles (right) along the AV axis by averaging the total Fgf8a concentration in each *xz*-plane. The model has the same orientation as the embryo in a).

**Figure 4.**
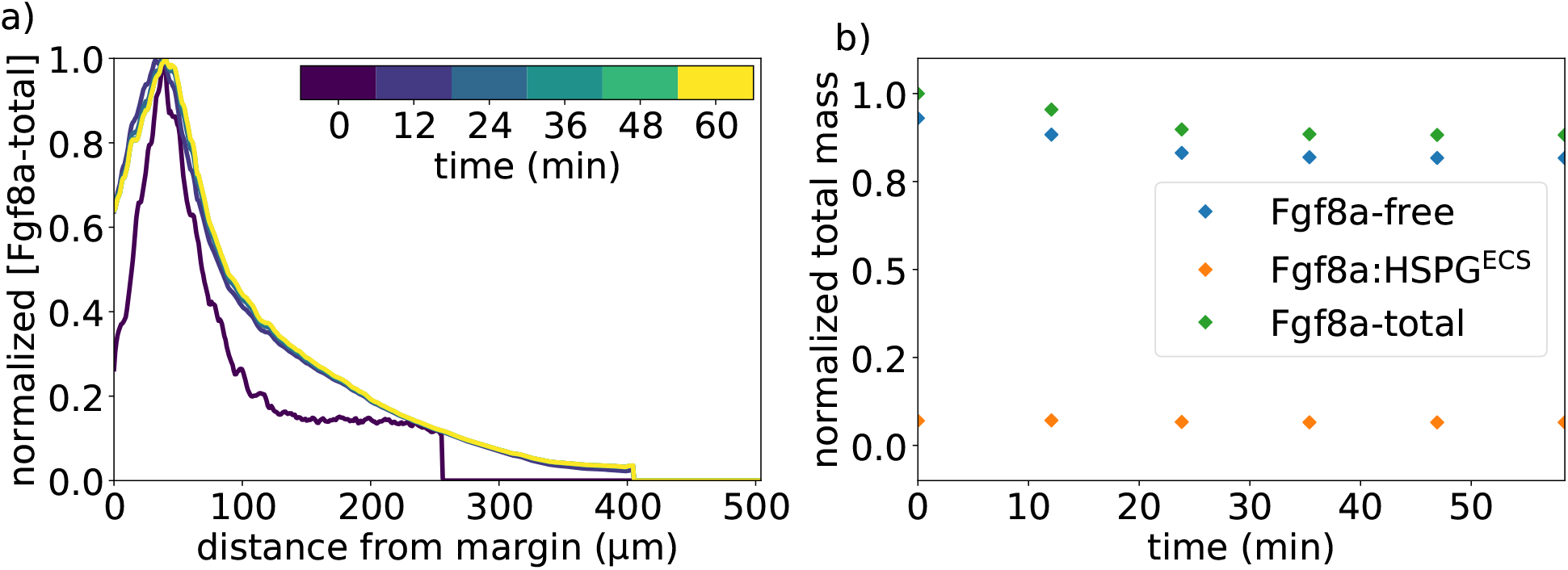
Simulated Fgf8a gradient using optimized parameters (Tab. 1) with experimental profile as initial condition. (a) Normalized AV concentration profiles of Fgf8a-total at different simulated times (color bar). All profiles are normalized by their respective maximum *xz*-plane-averaged concentration. The experimental profile used as initial condition is shown in dark purple (time 0). Simulated profiles reach a steady state within 12 min. (b) Normalized total mass of two Fgf8a fractions Fgf8a : HSPG^ECS^ and Fgf8a-free, along with their sum (Fgf8a-total). This set of parameters maintains a constant total Fgf8a mass and preserves the *in vivo* Fgf8a / Fgf8a : HSPG^ECS^ ratio of 93% / 7%).

Although maintaining the Fgf8a / Fgf8a : HSPG^ECS^ ratio and the overall profile, the simulated profile converges within 12 min to a steady state profile slightly flatter with a wider range than the experimental profile (Fig. 4a) and a lower peak concentration (see absolute concentration profiles in Suppl. Fig. S7). Possible explanations are that some of the parameter values is not correct, the experimental input profile is not at a steady state, or there is lower geometrical hindrance due to overestimated ECS connectivity and volume as a consequence of image-segmentation errors discussed in Section 2.5 and quantified in Section 3.2. Despite not perfectly maintaining the input profile’s slope and range, the simulated profile maintains a drop close to the margin, likely caused by a zonation effect in the marginal region with low cell density and poor connectivity to the rest of the ECS (cf. Fig. 3b). Furthermore, the normalized profiles maintain the overall shape of the *in vivo* gradient (Fig. 4a).

Overall, the parameters shown in Tab. 1 preserve the experimentally measured gradient at 60% epiboly and reproduce the *in vivo* ratio of Fgf8a / Fgf8a : HSPG^ECS^. We use these parameters throughout the manuscript.

## 3 Results

We reconstruct realistic 3D ECS geometries based on light-sheet microscopy volumes of zebrafish epiboly as described in Section 2.5. Exemplary optical sections of the light-sheet microscopy video are shown in Fig. 5. A full 3D visualization of the image-based model is shown in Fig. 5d – g for the time point at *≈*60% epiboly. We use this image-based model, simulation algorithm, and the parameter values described in Section 2 to study the emergence, maintenance, and robustness of the Fgf8a gradient in realistic ECS geometries of zebrafish embryos during epiboly. The computational model allows us to disentangle the influences of different molecular parameters and of the ECS geometry and shape. This enables testing for the sufficiency of an SDD mechanism of gradient formation and quantifying the sensitivity of the gradient to changes in the ECS geometry.

### 3.1 An SDD+HSPG-binding mechanism is sufficient to generate *de novo* Fgf8a gradients

We simulate *de novo* Fgf8a gradient formation in reconstructed zebrafish ECS geometries at *≈*40, 60, and 75% epiboly. The simulations start from initial concentrations [Fgf8a] = [Fgf8a : HSPG^ECS^] = [Fgf8a : HSPG^cellSurf^ ] = 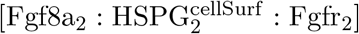 = 0 everywhere in the embryo to test whether an SDD+HSPG-binding mechanism is sufficient for the spontaneous emergence of an AV gradient.

The resulting concentration profiles and total mass over time at 60% epiboly are shown in Fig. 6a and b), respectively. A steady state is reached at *≈*30 minutes. The Fgf8a / Fgf8a : HSPG^ECS^ ratio remains almost constant (92.6% / 7.4%) throughout the simulated 60 minutes and very close to the *in vivo* measured ratio (93% / 7%). Interestingly the steady-state profile reached in the *de novo* simulation is similar to the steady-state profile when initializing with the experimental profile (Fig. 4), indicating an insensitivity of the steady state towards the initial condition.

**Figure 5.**
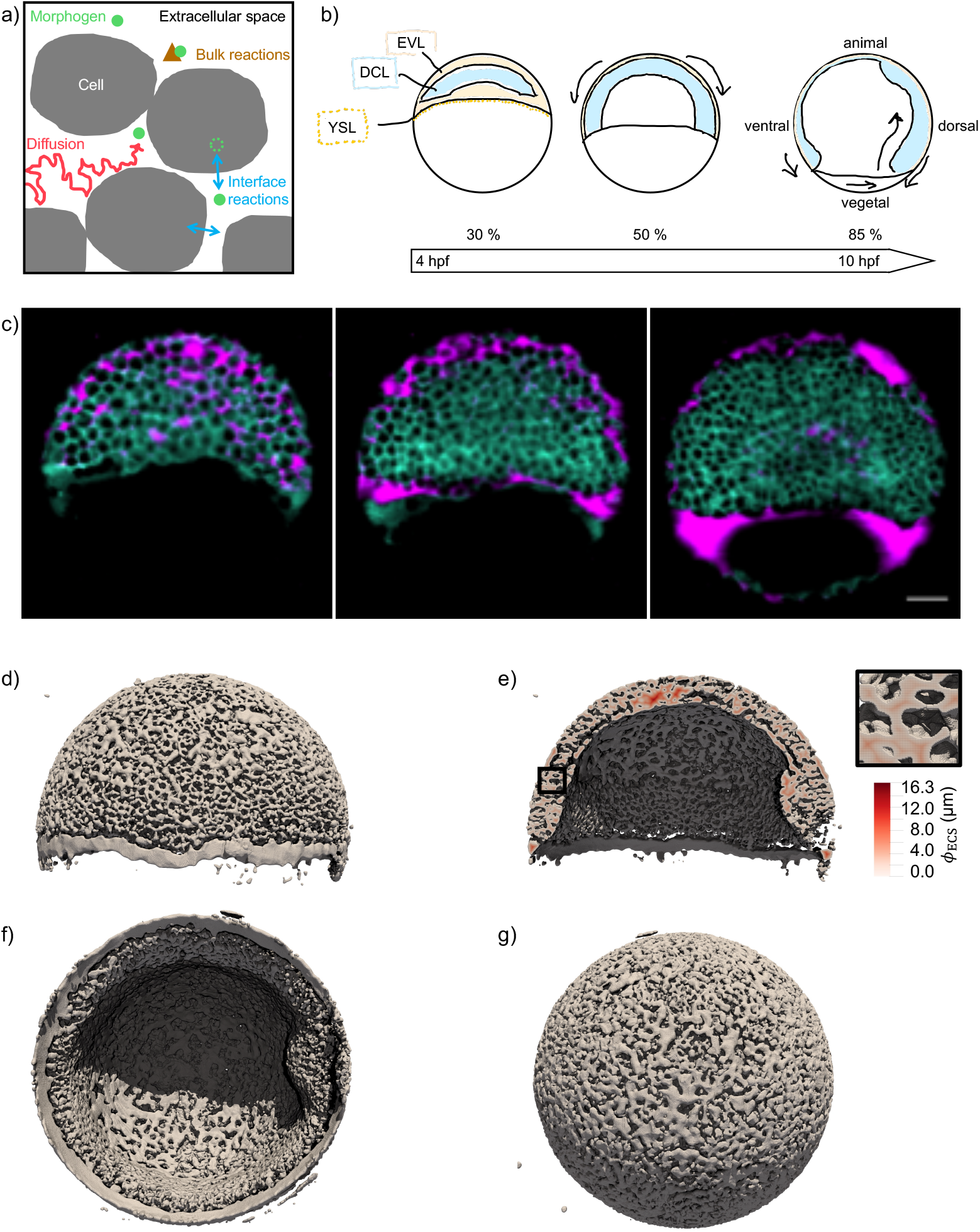
The extracellular space resembles an anisotropic porous medium during zebrafish epiboly. a) Schematic representation of diffusion in a porous medium with bulk and surface reactions. The tortuosity and imperfect connectivity of the pore-scale geometry lower the effective diffusivity and compartmentalize the space. Interfacial reactions depend on pore-surface area and pore accessibility. b) Illustration of zebrafish epiboly. Representative stages at around 40, 50, and 85% epiboly with deep-cell layer (DCL, blue), enveloping layer (EVL, yellow), and yolk syncytial layer (YSL, yellow dots). Starting at around 4 hpf (40% epiboly), the yolk moves toward the animal pole, and the blastoderm, composed of EVL, DCL, and YSL, starts spreading over the yolk. Toward the end of epiboly, the blastoderm almost completely engulfs the yolk, and asymmetry along the dorsal-ventral axis emerges. c) Light-sheet microscopy images of zebrafish epiboly. From left to right: exemplary optical sections from a Tg(bactin:hRas-EGFP) embryo at late blastula, early gastrula, and mid-gastrula stages, respectively, following image acquisition using a light-sheet fluorescence microscope and multi-view reconstruction (see Section 2.4). hRas-EGFP labels the cell membranes (green). ECS is marked by TMR-Dextran injection (magenta). Unidirectional views and single optical sections are shown for simplicity; scale bar: 50 µm. d) – g) Visualization of the SDF representation of the ECS surface at *≈*60% epiboly. We show an iso-surface rendering of *ϕ*_ECS_ = 0 in gray. d) Side view oriented as in b) and c). e) Cross-cut view with *ϕ*_ECS_ values in increasing distance to the ECS surface shown in shades of red (color bar). Face-on views from vegetal (f) and animal pole (g).

**Figure 6.**
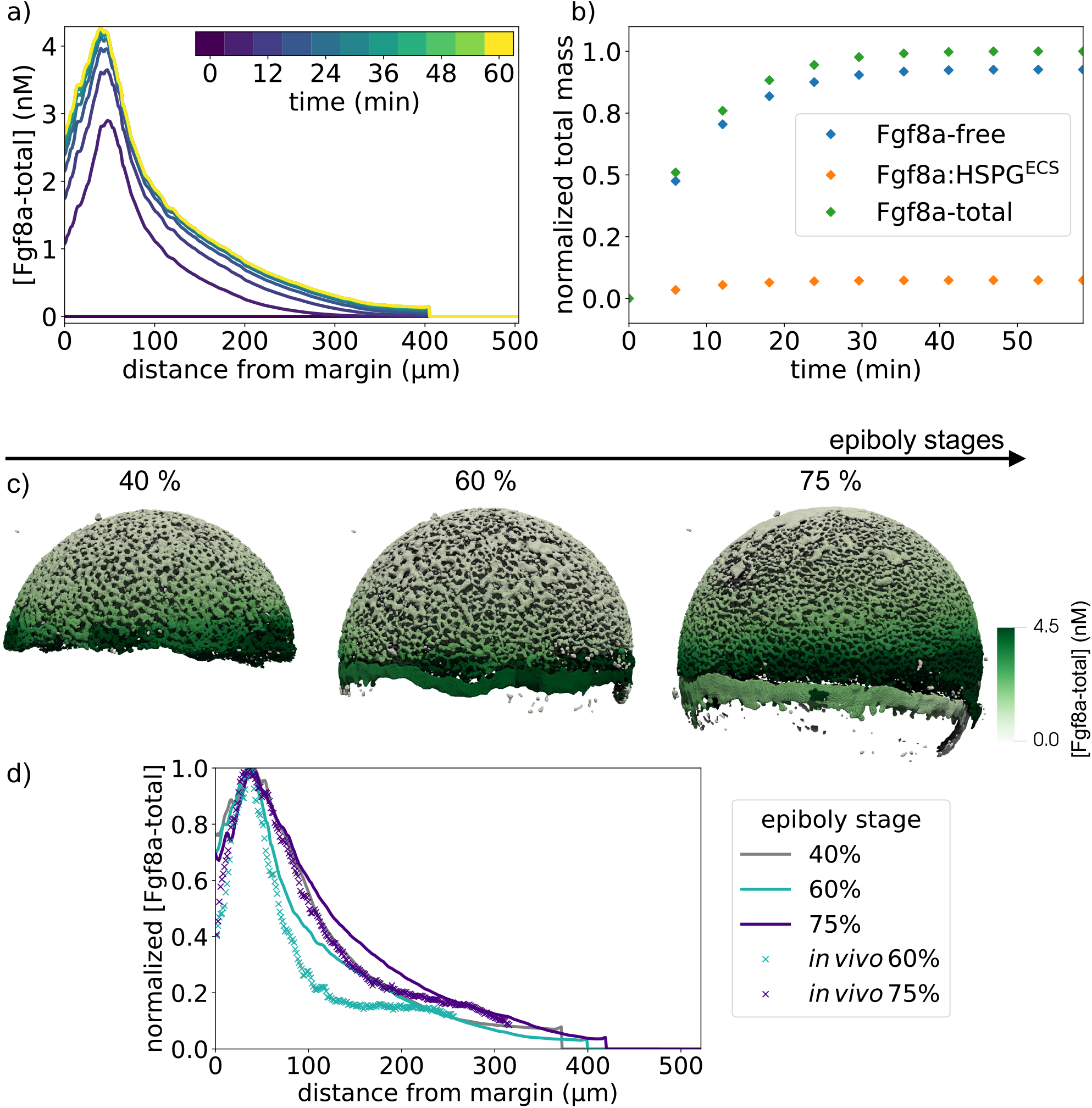
An SDD+HSPG mechanism is sufficient to explain the spontaneous emergence of the Fgf8a gradient in realistic ECS geometries. All Fgf8a concentrations in the simulation were initially set to zero throughout the embryo. a) Evolution of the emerging AV gradient profile of [Fgf8a − total] over time (color bar). The gradient reaches a steady state at *≈*30 min. b) The Fgf8a-free / Fgf8a : HSPG^ECS^ ratio remains almost constant (92.6% / 7.4%) and close to the experimentally measured values (93% / 7%) throughout the simulated 60 minutes. c) Visualization of the simulated Fgf8a-total concentrations in the ECS geometries at 40, 60, and 75 % epiboly. d) Simulated (lines) and experimentally measured (symbols) AV profiles at different stages (colors, inset legend). Experimental profiles were only available at 60 and 75% epiboly. Simulated profiles are computed at *t* = 60 min.

**Figure 7.**
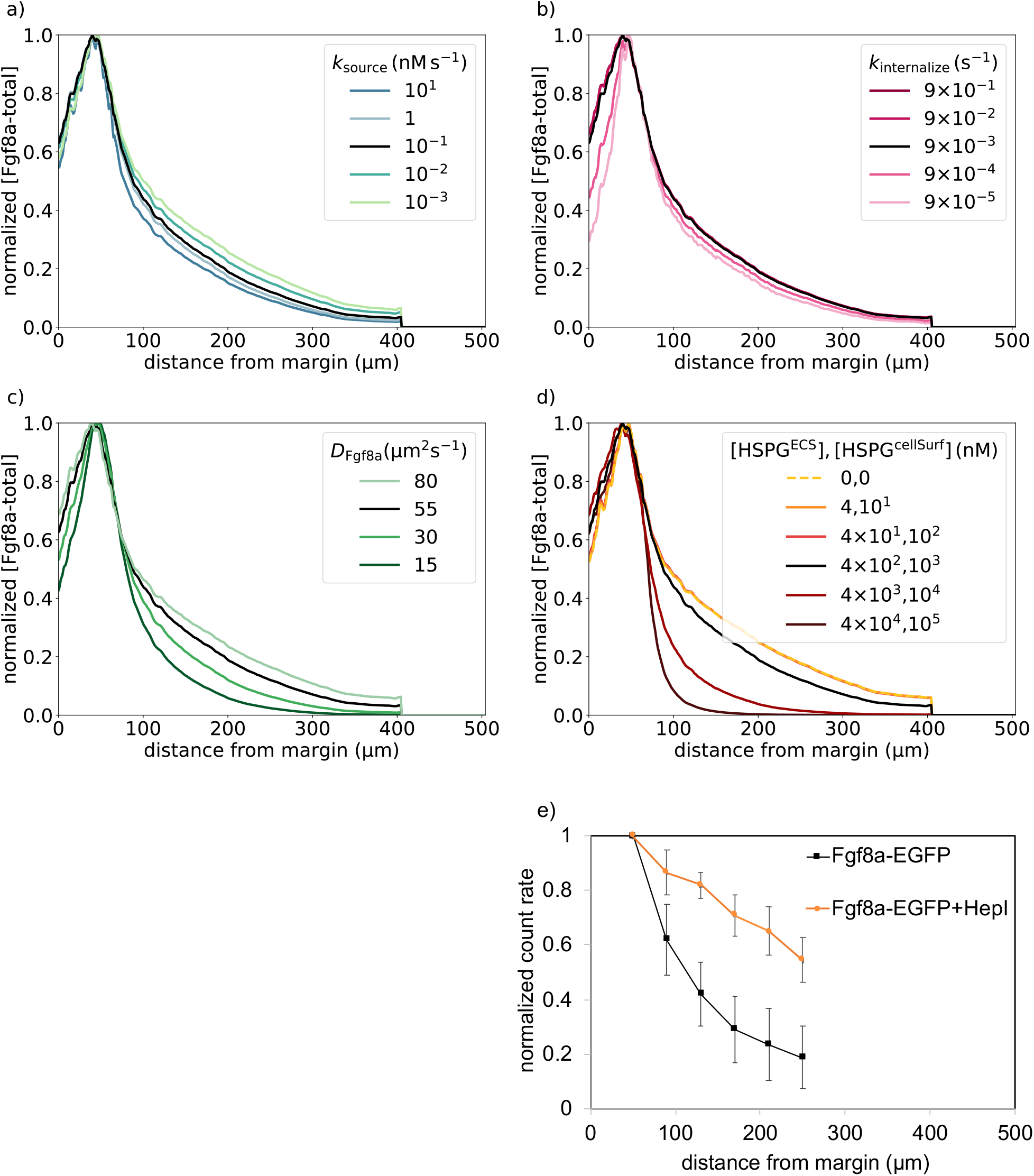
Sensitivity of simulated steady-state Fgf8a gradients to variations in the model parameters a) – d) and *in vivo* gradient sensitivity to HepI injection e). The simulated gradient shows remarkable robustness against changes in source (a) and sink rates (b) over four orders of magnitude (color, inset legend). Changes in the effective diffusivity, by changing either the molecular diffusion coefficient of Fgf8a (c) or HSPG concentrations (d), strongly affect the gradient. The higher HSPG binding and the smaller the diffusion coefficient, the steeper and shorter the gradient. The baseline gradient (Section 3.1) is shown as a solid black line in all panels. All profiles are normalized by their respective maximum *xz*-plane-averaged concentration. Profiles are shown at *t* = 60 min. e) FCS count rates of Fgf8a without (black squares) and with (orange circles) HepI injection. HepI cleaves off the HS side-chain of HSPGs. The Fgf8a gradient of HepI-injected embryos is shallower. Figure reproduced with permission from Harish et al. (2023). Data are represented as mean *±* standard deviation. 23

Overall, the simulated steady-state profile is similar to the experimental profile (Fig. 3a) but shows a slightly flatter gradient with longer range. A possible reason for this deviation is that the *in vivo* profile has been averaged over a sub-volume of the embryo (yellow dotted lines in Fig. 3a), whereas the simulated profile is computed across the entire ECS. Another reason could be modeling errors in the ECS geometry reconstruction or the simulation parameters. Indeed, the image segmentation tends to overestimate ECS thickness, reducing geometrical hindrance in the model compared to the real embryo and enabling Fgf8a to diffuse faster and further, which would explain the gradient flattening and increased decay length.

We repeat the *de novo* gradient formation simulation in ECS geometries t *≈*40 and *≈*75 % epiboly. The results are shown in Fig. 6c and d. Again, the simulated profiles are slightly flatter than the experimentally measured ones but are overall similar and show the same trend over developmental stages. Also, in the model, propagation lengths increase as epiboly progresses, probably due to a widening of the source band.

Interestingly, in addition to the AV gradient, the simulation also reproduces the DV gradient observed *in vivo* at 75% epiboly (see Supplementary Note S2, Suppl. Figs. S2 and S3). Since, in the model, the rates of all sources, sinks, and HSPG binding are invariant along the DV axis, this gradient can be interpreted as a consequence of ECS geometry asymmetry. In addition to this sufficient geometric asymmetry, earlier patterning events have been proposed to contribute to DV gradient formation [44, 97].

Taken together, these results show that the SDD+HSPG-binding mechanism modeled here is sufficient to cause the spontaneous emergence of AV and DV gradients in zebrafish embryo ECS geometries during epiboly. The mechanism also maintains the experimentally measured Fgf8a population fractions and explains the gradient evolution across epiboly stages.

### 3.2 Fgf8a normalized gradients are robust to changes in rate constants but sensitive to changes in ECS geometry

Factors such as temperature and DNA distribution can cause fluctuations in gene-expression levels and therefore reaction rates. Gradient robustness is the capacity to buffer the effect of such fluctuations and is important to ensure that tissues develop correctly over a range of environmental conditions. Using our model in ECS geometries at 60% epiboly, we quantify how the rates of sources, sinks, HSPG binding, and diffusion influence the gradient profile. In addition, we study the impact of ECS geometry on the gradient.

#### The normalized gradient profile is robust against changes in source and sink rates

Morphogen gradients are surprisingly robust against changes in the source rate [35, 36, 76, 134]. In previous models, this robustness has been explained by nonlinearities of the ligand current when the morphogen is transported by transcytosis [16] and nonlinear degradation in the free diffusion model [37].

To test whether nonlinearities due to the complex ECS geometry and spatially heterogeneous degradation of our model are sufficient to reproduce this robustness, we simulate *de-novo* Fgf8a gradient formation for different *k*_source_, keeping all other parameters fixed, until a steady state is reached at *t*_max_ = 60 min. The resulting AV gradient profiles in Supplementary Fig. S8a show that higher *k*_source_ lead to higher overall concentration levels, an expected consequence of increased morphogen secretion. However, normalizing by each profile’s maximum concentration, as also done in experimental measurements, reveals that the gradient shape is almost invariant across 4 orders of magnitude in *k*_source_ (see Fig. 7a. Compared to the baseline *k*_source_ = 0.1 nM s^−1^, the gradient becomes only slightly flatter when decreasing *k*_source_ and slightly steeper when increasing *k*_source_. The deviations are, however, very small.

This shows that the mechanisms modeled here are sufficient to explain the experimentally observed gradient robustness [35, 36, 76, 134]. In our simulations, the normalized Fgf8a AV gradient is robust to source-rate changes across 4 orders of magnitude, from 10^−3^ … 10. A possible explanation for this robustness is that the sink term scales with the morphogen concentration, thereby buffering the effects of variations in the production.

For the sink rate, *in-vivo* observations suggest that the gradient is not robust. It has been shown that modulating Fgf8a endocytosis in live zebrafish embryos makes normalized gradients shallower when inhibiting endocytosis and steeper when upregulating endocytosis [133]. We test the behavior of our model by changing *k*_internalize_ while keeping all other parameters unchanged. The resulting gradient profiles in Fig. S8b suggest that decreasing *k*_internalize_ leads to higher concentration levels, which is an expected consequence of reduced morphogen degradation. Normalizing the profiles, however, it can again be seen in Fig. 7b that the gradient shape is almost unaffected by changing *k*_internalize_ across 4 orders of magnitude. A significant difference can only be seen within 50 µm from the margin, where the gradient becomes steeper for smaller *k*_internalize_ than the baseline *k*_internalize_ = 9· 10^−3^ s^−1^ but only slightly flatter for *k*_internalize_ larger than baseline.

The simulation does therefore not explain the experimentally observed sensitivity of the gradient to endocytosis rates. The gradient flattening observed *in vivo* is therefore not caused by the molecular mechanisms modeled here. Instead, we speculate that it could be caused by side effects of a dominant-negative mutation of Dynamin, a GTPase that pinches off endocytic vesicles from the plasma membrane. Changes in Dynamin function are therefore not specific to Fgf8a endocytosis and may affect other feedback loops and downstream signaling cascades, too.

In conclusion, the simulation results show that an SDD+HSPG mechanism is sufficient for generating gradients whose normalized profile is robust to changes in *k*_source_ and *k*_internalize_ across 4 orders of magnitude. This robustness does, however, not transfer to the absolute concentration gradient whose overall levels increase for increasing *k*_source_ and decreasing *k*_internalize_.

#### Slower diffusion leads to steeper and shorter gradients

Next, we quantify gradient robustness against changes in morphogen diffusion. We simulate *de-novo* gradient formation using different diffusion coefficients of Fgf8a-free. The results are shown in Figs. S8c (absolute values) and 7c (normalized). Compared to the baseline diffusivity of *D*_Fgf8a_ = 55 µm^2^s^−1^, lower diffusion coefficients result in steeper gradients and shorter propagation lengths with narrower, higher peaks at the source. Increasing the diffusion coefficient leads to gradient flattening, longer propagation, and lower peak concentration.

This shows that the steady-state profile of the morphogen gradient depends on the morphogen diffusion coefficient. The dependence is much stronger than for the source and sink rates, where a 10,000-fold variation led to smaller changes in the gradient than a five-fold change of the diffusion constant. Faster diffusing morphogens can propagate further and have longer signaling ranges in the embryo, whereas slower diffusing morphogens maintain higher concentrations near the source and act more locally. This suggests that gradient formation could more generally be influenced by any factor hindering or promoting morphogen diffusion. This includes the geometric tortuosity of the diffusion space and HSPG binding.

To test the influence of HSPG binding on gradient formation, we simulate *de-novo* gradient formation for different [HSPG^ECS^] and [HSPG^cellSurf^ ]. As shown in Fig. 7d, higher HSPG concentrations lead to gradients that are considerably steeper with shorter propagation lengths. Decreasing HSPG concentrations 10-fold below the baseline flattens the gradient. HSPG therefore acts on the gradient by modulating the effective diffusivity through transient matrix binding.

Flatter gradients for lower HSPG concentrations are in agreement with *in vivo* HepI-injection experiments. HepI is an enzyme that cleaves the HS side chains of HSPG, reducing the slow-diffusing Fgf8a fraction, thereby causing a shallower gradient [50, 133] as measured by FCS and shown in Fig. 7e. HepI injection also increased the overall Fgf8a concentration in the ECS, which has been explained by Fgf8a dissociating from cell-surface HSPGs [50]. Our simulations reproduce this observation, showing higher Fgf8a peak concentrations at lower HSPG concentrations (see Fig. S8d).

Interestingly, decreasing HSPG concentrations below [HSPG^ECS^] = 40 nM and [HSPG^cellSurf^ ] = 100 nM does not further flatten the gradient. Even when setting all HSPG concentrations to zero and modeling gradient formation by an SDD mechanism, the same profile emerges as for [HSPG^ECS^] = 40 nM and [HSPG^cellSurf^ ] = 100 nM (see coinciding lines in Fig. 7d). This means that a minimal HSPG concentration is required to have a relevant effect on the gradient. This makes sense as below the baseline, the freely diffusing Fgf8a fraction constitutes *>* 92 % of the total Fgf8a, dominating the gradient profile. This also shows that under the current modeling assumptions, an SDD mechanism is sufficient for the spontaneous formation and maintenance of some gradient, which is, however, flatter than the baseline and the *in vivo* gradient.

Both *in vivo* and *in silico*, HSPG^cellSurf^ is involved in receptor binding and endocytosis of Fgf8a. Decreasing [HSPG^cellSurf^ ] thus also decreases sink fluxes. While changing the sink rate *k*_internalize_ did not affect the simulated gradient profile (Fig. 7), reducing the effective sink flux does.

Concluding, the present results show that morphogen gradients are sensitive to changes in morphogen diffusivity. Both reducing the molecular diffusion coefficient and the diffusive hindrance by HSPG binding have a similar effect, leading to steeper gradients with shorter range and higher peak near the source. This is in agreement with previous 1D models showing that lower diffusion coefficients lead to steeper and shorter gradients [125] and with *in vivo* experiments showing that HSPG binding is crucial for maintaining high concentrations near the source and for restricting gradient propagation lengths [50].

#### Tortuous ECS geometries stabilize Fgf8a gradients

The effect of decreasing HSPG concentrations is much smaller than when increasing them (Fig. 7d). Indeed, the gradient remains almost unchanged when reducing [HSPG^ECS^] or [HSPG^cellSurf^ ] by more than 10-fold. This suggests that the diffusive hindrance from HSPG binding is dominated by the tortuosity of the ECS geometry. This is consistent with HSPG being mostly localized near cell surfaces, mirroring the ECS geometry.

We use our realistic 3D ECS geometry model to quantify the effect of diffusive tortuosity on Fgf8a gradients. For this, we perturb the modeled ECS geometry and quantify the influence of these perturbations on simulated Fgf8a gradients. Such perturbations are straightforward in the numerical description of the ECS as a signed-distance function. Changing the surface level-set from the standard *ϕ*_ECS_ = 0 to other values uniformly changes the ECS tube diameters and thus the ECS density, connectivity, and volume.

We vary the geometry in steps of *h* from *ϕ*_ECS_ = 3*h* (thinner ECS, higher tortuosity) down to *ϕ*_ECS_ = −1*h* (thicker ECS, lower tortuosity) in comparison with the baseline *ϕ*_ECS_ = 0. The unit *h* is the grid spacing, i.e., twice the pixel size of the light-sheet microscopy images. Examples of resulting perturbed geometries are shown in Fig. 8a for *ϕ*_ECS_ = −1*h* and *ϕ*_ECS_ = 3*h*. For these perturbed geometries, we compute steady-state Fgf8a gradient profiles, keeping all model parameters at their default values as given in Table 1.

**Figure 8.**
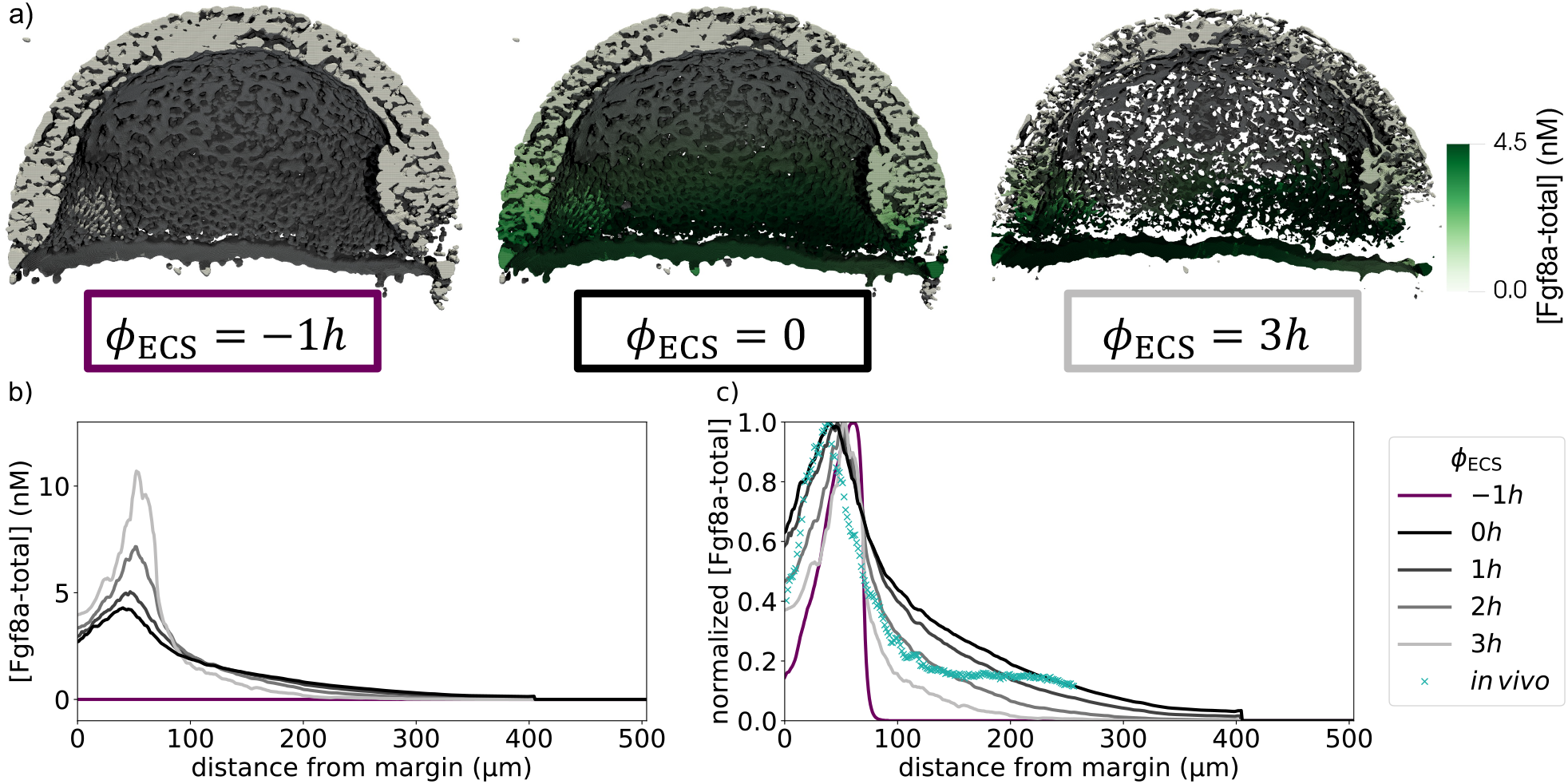
ECS geometry controls Fgf8a gradient steepness and range. To test how ECS tortuosity influences Fgf8a gradients, we simulated *de-novo* gradient formation for different ECS geometries. a) Clipped visualizations of the baseline ECS geometry (center) and two perturbed versions (left, right) with simulated Fgf8a concentrations overlaid in green, scaled to [Fgf8a − total]_max_ = 4.5 nM (color bar). The ECS tube thickness is perturbed by changing the boundary location along *ϕ*_ECS_, i.e., shifting down by one grid spacing *h* (2 pixels) to obtain thicker tubes and increasing by 3*h* to obtain thinner tubes. b) Absolute AV concentration profiles at *t* = 60 minutes. Colors correspond to different geometries (inset legend). Compared to the baseline geometry (solid black line), decreasing ECS tube sizes limits Fgf8a propagation lengths and leads to a steeper gradient with a higher peak (lines shades of gray). Increasing the tube sizes and imposing sources and sinks only at the surfaces of the remaining cells causes the gradient to vanish (violet line). c) The same profiles normalized to their respective maximum concentration values. The *in vivo* profile at 60 % epiboly (symbol) is more similar to the simulated profile for thinner tubes of *ϕ*_ECS_ = 2*h*.

The resulting absolute concentration profiles in Fig. 8b show that the thinner the ECS tubes, the more elevated the peak near the source, the steeper the gradients, and the shorter the range. The steeper, shorter gradients for higher ECS tortuosity are also confirmed in the normalized profiles (Fig. 8c). This shows that morphogen gradients are sensitive to ECS geometry and that geometric hindrance is sufficient to tune morphogen propagation lengths.

Thickening the ECS tubes by only 1*h* leads to negligible Fgf8a concentrations (*<* 0.007 nM) and a flat profile, shown in Fig. 8b. In the normalized profile, a peak at the source becomes apparent with a small peak concentration of 0.001 nM. This amounts to *≈*10 times the secreted concentration per simulation time step, showing that this peak can likely be attributed to instantly secreted Fgf8a, partly immobilized at the cell surface of secretion as Fgf8a : HSPG^cellSurf^ and 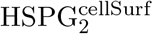: Fgfr_2_. Away from the immediate source region, concentrations are zero and no gradient exists.

The absence of a gradient, in this case, can be explained as follows: The lower ECS tortuosity enables the morphogen to quickly diffuse away from the source and distribute throughout the target tissue, where it is degraded. In addition, the effective source flux is reduced as the surface area between ECS and deep cells is reduced by thickening the tubes. This is not balanced by the—for the same reason—lower effective sink flux. This shows that under the current modeling assumption of restricting the source to the deep-cell surfaces, increasing ECS thickness by 1*h* compared to the baseline results in a tortuosity insufficient to generate and maintain a gradient.

Taken together, these results show that the tortuous ECS geometry plays an important role in establishing and maintaining robust morphogen gradients. The geometric tortuosity regulates source peak height, gradient steepness, and propagation length. In geometries of insufficient tortuosity, gradients is not maintained. Regulating gradient formation and maintenance, therefore, involves adjusting extracellular HSPG concentrations and ECS tortuosity to the morphogen diffusion coefficient. The embryo can thus control the morphogen gradient by tuning HSPG secretion and the shape of the interstitial space, e.g., by regulating cell adhesion and tissue rigidity [80, 92]. The present results suggest that the ECS tortuosity is tuned to just the minimum required to support long-range gradients, as already reducing it by one grid unit causes the gradient to vanish.

### 3.3 Porous-media characteristics of the ECS regulate gradient shape

So far, the results show that Fgf8a AV gradients are robust to changes in source and sink rates when simulated in realistic ECS geometries but sensitive to changes in the ECS geometry. Since the gradients are also sensitive to changes in the molecular diffusion constant and in transient HSPG binding, this suggests that diffusive hindrance may be a core mechanism in morphogen gradient emergence, maintenance, and robustness. Diffusive hindrance is a core concept of transport in porous media [3, 10, 14, 34, 38, 45, 100, 105, 106, 112, 123]. In the present system, diffusive hindrance results from the joint effect of geometric tortuosity, transient HSPG binding in the ECS and at the cell surfaces, receptor complex formation, and the finite molecular diffusion constant of the morphogen. In this sense, the ECS geometry, with its extracellular HSPG, effectively acts as a porous medium regulating Fgf8a gradient shape.

Our model resolves the porous medium explicitly by resolving geometry and binding reactions at the “pore scale”. To test whether Fgf8a gradient shape can be explained by an effective porous medium, we compare simulations in a fully resolved ECS geometry with simulations in an upscaled homogenization [19, 54, 68]. In this homogenization, the geometry is simplified to a spherical shell by extracting the outer surfaces of our ECS geometry model at 60% epiboly, filled with a uniform porous material (see Fig. 9). This porous material models the coarse-grained effect of ECS geometry and diffusive hindrance without explicitly resolving them. In this simplified model, surface reactions and diffusive tortuosity are represented by uniformly distributed effective volume rates. The source is uniformly distributed within a deep-cell band at the blastoderm margin with the same width as in the fully resolved geometry model (70 µm), as shown in Fig. 9a. Receptor-mediated endocytosis is modeled throughout the DCL, as shown in Fig. 9b.

**Figure 9.**
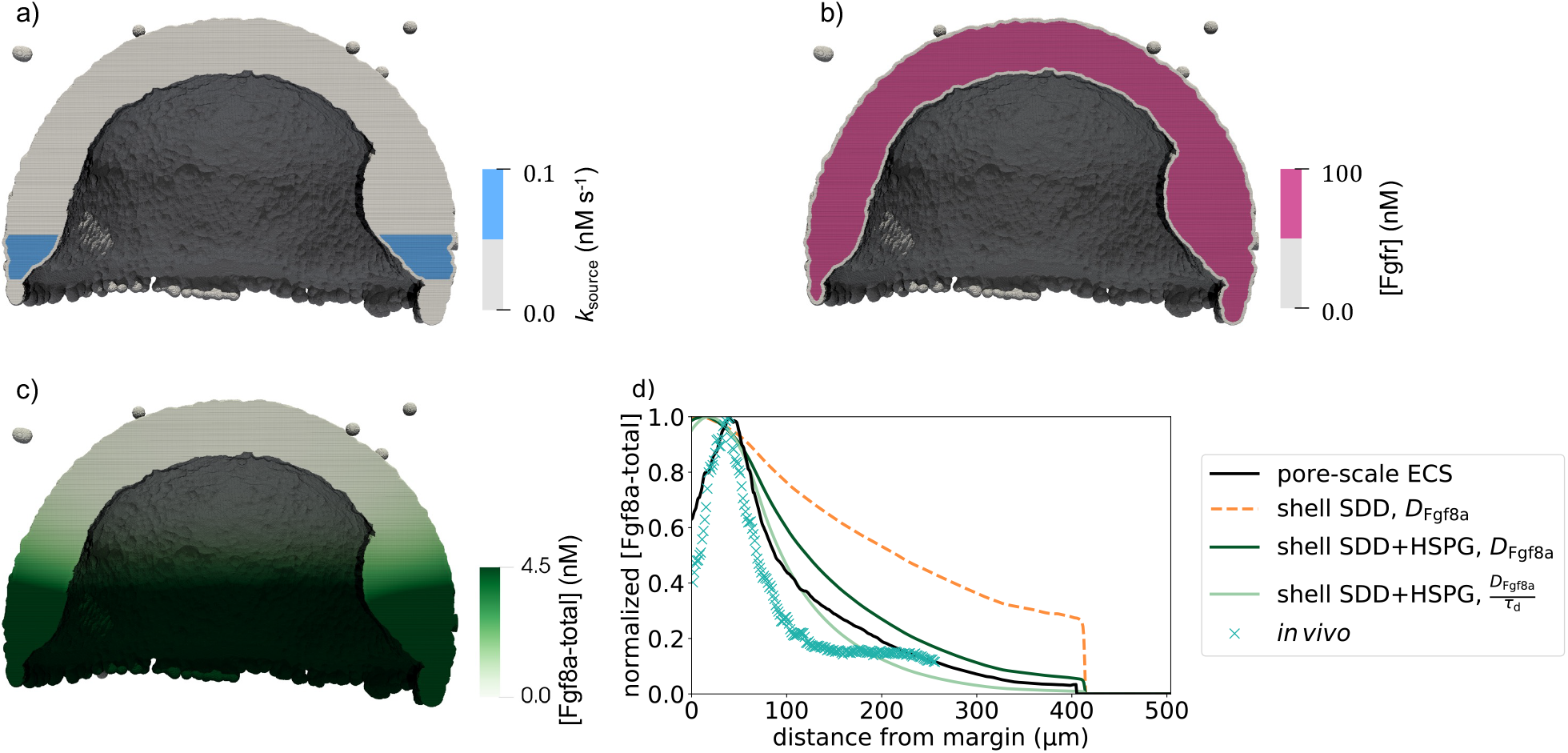
Porous-media characteristics of ECS can be partly upscaled in a shell geometry. Clipped simulation visualization of (a) the source in a 70 µm DCL band at the margin, (b) the uniformly distributed sink ([Fgfr]) in the DCL, and (c) the Fgf8a gradient at *t* = 60 minutes (concentration color bar) using the parameters from Table 1. (d) Modeling the ECS geometry as a spherical shell without HSPG leads to a longer, flatter gradient (orange dashed line) compared to the baseline of the pore-scale model (black line). Gradient steepness and range are partly recovered by upscaling HSPG binding by uniformly distributed HSPG^ECS^ and HSPG^cellSurf^, and it is fully recovered when further upscaling the tortuosity by an effective diffusion coefficient using *τ*_d_ (as defined in Supplementary Note S5). This does, however, not apply to local geometry effects such as the zonation near the source as found in the *in vivo* profile (symbol) and reproduced by the pore-scale model. AV profiles are reported at *t* = 60.0 min.

We first quantify the effect of HSPG binding by simulating Fgf8a gradient formation by an SDD mechanism without HSPG binding and Fgf8a degradation at rate *k*_decay_ in the entire shell. Using this homogenized model, we simulate *de-novo* gradient formation with the parameters given in Table 1 until a steady state is reached at *t* = 60 min. The normalized gradient profiles in Fig. 9d show that the gradient in the spherical shell geometry emerging from an SDD mechanism without HSPG (orange dashed line) does not reproduce the baseline gradient (black line). The gradient in the spherical shell is shallower with a longer propagation length and lacks the zonation near the margin. The latter is because in the coarse-grained model, the marginal region is fused with the shell, whereas in the real geometry, the margin is only weakly connected to the rest of the ECS. The flatness and long range can be explained by missing diffusive hindrance from HSPG and Fgfr binding and ECS tortuosity.

To upscale diffusive hindrance by HSPG binding and complex formation, we assume uniformly distributed HSPG^ECS^ and HSPG^cellSurf^ in the entire shell and receptor-mediated endocytosis throughout the DCL (see Fig. 9b). The diffusive hindrance provided by HSPG binding and receptor-complex formation in the entire volume of this geometry is stronger than in the baseline ECS geometry, where these effects are restricted to the cell surfaces. Figure 9d shows that under these modeling assumptions, *de-novo* formed gradients are steeper and shorter (dark-green line) than with an SDD mechanism alone. Although this gradient is still shallower than the baseline and lacks the zonation near the source, gradient steepness is partly recovered by this upscaling of HSPG binding.

In addition to HSPG binding, the effect of geometric hindrance can be upscaled using the diffusive tortuosity *τ*_d_ of the ECS geometry, as defined in Eq. (S1), Supplementary Note S5. Using the fully resolved ECS geometry model, we numerically compute *τ*_d_ = 2.5 on average across the entire ECS, as detailed in Supplementary Note S5. This factor describes how strongly the ECS geometry alone hinders Fgf8a diffusion. Limited by the light diffraction limit and segmentation errors, our geometry model likely overestimated the thickness of ECS structures due to light. We, therefore, expect the real *τ*_d_ to be even larger [135].

Simulating gradient formation in the spherical shell with upscaled diffusion coefficient 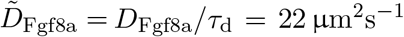 indeed recovers gradient steepness and range (see Fig. 8d, light-green line). The geometric zonation effect near the source, however, is still not reproduced, as this local effect is not captured by the averaged tissue-wide factor.

In summary, these results show that the ECS effectively acts as a porous medium when modulating morphogen gradient shape. In this, geometric tortuosity acts in concert with matrix and receptor binding to create the required diffusive hindrance. Hindrance from matrix binding alone does not account for the correct gradient slope. The effective geometric hindrance, as represented by the upscaled tortuosity, restores the global gradient shape apart from local zonation effects. This shows that the porous-media characteristics of the ECS geometry are sufficient to explain morphogen gradient shape.

## 4 Discussion

Combining image-based three-dimensional geometry modeling with geometry-adaptive numerical simulation methods enabled reconstituting morphogen gradient emergence and maintenance *in silico* in realistic embryo geometries with fully resolved interstitial space, surface, and bulk biochemistry. We reconstructed pore-scale extracellular space (ECS) geometries of zebrafish embryos during epiboly from light-sheet microscopy videos and numerically solved the governing equations of Fgf8a gradient formation in those geometries. The simulations were validated against *in vivo* fluorescence intensity profiles and single-molecule fluorescence correlation spectroscopy (FCS) measurements, showing that the *in vivo* gradient can be explained by a source-diffusion-degradation (SDD) mechanism with additional heparan-sulfate proteoglycan (HSPG) binding in the extracellular matrix. We found that the Fgf8a gradient is robust against changes in the Fgf8a source and sink rates but sensitive to perturbations of the ECS geometry, suggesting that the geometry plays an important role in gradient shaping and maintenance. This indicates that the “SDD+HSPG+geometry” mechanism proposed here is sufficient to explain the emergence, scaling, maintenance, and robustness of morphogen gradients.

We showed that gradient steepness and range are regulated by diffusive hindrance through HSPG-binding and tortuous ECS geometries. The embryo can thus control gradient shape by tuning heparan-sulfate affinity and by modulating the HSPG secretion rate. In addition, ECS tortuosity can be controlled by modulating cell density and adhesion, varying the interstitial space. Modulation of cell adhesion can be observed, for example, during tissue fluidization of the central deep cells at the animal pole during the onset of doming [80, 92]. The aggregate diffusive hindrance is key to preserving high morphogen concentrations near the source and regulating gradient steepness and range.

Diffusive hindrance is a core characteristic of porous media. We thus suggest that the ECS in the developing embryo effectively acts as a porous medium toward the morphogen gradient. Indeed, we were able to restore the long-range gradient profile by coarse-graining (i.e., computational upscaling) the average porous-media characteristics of the ECS geometry into a spherical shell. Coarse-grained average porous media characteristics, however, were not able to account for the zonation effect near the blastoderm margin, where reduced ECS pore connectivity leads to an offset concentration peak. This effect was reproduced in the fully resolved realistic ECS geometry, whereas it was naturally lost in an average upscaling. This means that although effective porous media characteristics of the ECS govern the long-range gradient shape and range, local features may still intricately depend on the geometric details of the diffusion space.

This dependence on local geometric detail also implies that our results are sensitive to imaging, image-segmentation, and geometry-modeling errors. Although the image-based geometry used here was sufficiently realistic to demonstrate the pore-scale effects of geometry and explain the experimental observation, it likely underestimated real ECS tortuosity. The model could be refined in the future by enhancing image acquisition and segmentation.

The present model also neglects flows in the ECS due to tissue growth and cell motion, assuming that diffusive transport in the ECS is significantly faster than the time scale of tissue deformation. This is an approximation, as the real embryo geometry varies continuously over time. Although dilution effects are likely negligible during epiboly, as the ECS volume does not change (see Supplementary Note S4), cell movement has been proposed to contribute to gradient formation by additional advective transport [72]. Quantifying the contributions from advection and deformation would require explicitly modeling tissue dynamics [62, 93]. Tracking the ECS geometry over time to derive continuous deformation maps requires image data with a higher time resolution than the light-sheet microscopy video used here. Once available, such deformation maps could be included in our model by correspondingly evolving the level-set function [12] and simulating the induced flows in the ECS [111]. Such simulations would also enable studying how morphogen gradients scale across different tissue sizes during growth [23, 43, 127].

While we focused on Fgf8a gradients in zebrafish embryos during epiboly, the presented modeling workflow and computer simulation algorithm are general and transferable to other types of morphogens and systems. Beyond the case of zebrafish epiboly, many morphogen gradients form and act in complex-shaped diffusion spaces involving surface reactions such as secretion and endocytosis [83]. We have shown here that the geometry of the diffusion space can play an important role in shaping and stabilizing a morphogen gradient, where the tissue thus acts as a porous medium whose physicochemical and geometric characteristics regulate gradient dynamics.

The regulation of biological reaction-diffusion processes by porous-media characteristics of the tissue may constitute a general principle. Such processes typically take place in complex geometric environments and are subject to surface and bulk reactions. Diffusive hindrance from binding reactions and geometric tortuosity, the two key mechanisms identified here, are thus likely universal. Apart from morphogen gradients, this applies to other diffusive transport processes in complex tissue geometries, including bone repair [74], wound healing [7], nutrient transport in the meniscus [119], tumor cell proliferation [40], and therapeutics diffusion through the brain ECS [128]. On the other hand, pathological conditions that cause cell swelling, e.g., during ischemia, increase the diffusive tortuosity, hindering the diffusion-mediated supply of those tissues [52]. These processes are — like morphogen gradient formation —modulated by the porous media characteristics specific to each tissue type, particularly the pore-scale tissue geometry.

## Data and code availability

Code: The simulation software used in this work is available under the 3-clause BSD license and free of charge from https://github.com/mosaic-group/openfpm. We provide the specific code of this paper on https://git.mpi-cbg.de/mosaic/reactiondiffusion_imagebased_porousmedia.git.

## Acknowledgments

We thank Dr. Nandu Gopan, Alejandra Foggia, and Dr. Johannes Pahlke (all Sbalzarini lab, CSBD Dresden), as well as past and present members of the Brand lab at CRTD Dresden, and Dr. Anastasia Solomatina (Leibniz-HKI Jena) for fruitful discussions and comments on the manuscript, and Dr. Pietro Incardona (University of Bonn) for help with the OpenFPM implementation of the simulation software. We thank Dr. Michele Marass (MPI-CBG, Dresden) for editorial advice. We thank Prof. Dr. Frank Jülicher (MPI-PKS, Dresden) for his comments on the manuscript.

## Author Contributions

Conceptualization, J.S., R.K.H, I.F.S., M.B.; Methodology, J.S., R.K.H, I.F.S., M.B.; Software, J.S.; Validation, J.S., R.K.H, I.F.S., M.B.; Formal Analysis, J.S., R.K.H; Investigation, J.S., R.K.H, I.F.S., M.B.; Resources, I.F.S., M.B.; Data Curation, J.S., R.K.H; Writing – Original Draft, J.S., R.K.H, I.F.S., M.B.; Writing – Review & Editing, J.S., R.K.H, I.F.S., M.B.; Visualization, J.S., R.K.H; Supervision, I.F.S., M.B.; Project Administration, I.F.S., M.B.; Funding Acquisition, I.F.S., M.B..

