## Supplementary Notes and Material for "Morphogen gradients are regulated by porous media characteristics of the developing tissue"

### S1 Grid initialization with experimental gradient

To determine parameters that maintain the *in vivo* gradient, we initialized concentration fields with the experimental intensity profile at 60% epiboly (Fig. 3a)), as this profile has more data points than the FCS count profile shown in Fig. 7e. The intensity profile is one-dimensional and provides average values at discrete locations along the AV axis. In our three-dimensional model, we distribute these values uniformly over $xz$-planes along the same axis after margin alignment, with a Fgf8a-free / Fgf8a : HSPG$^{\mathrm{ECS}}$ ratio of 93% : 7%. It has been shown that Fgf8a concentrations are higher at the cell membranes than in the ECS [49]; by which factor is, however, unknown, but the factor is not large. To nevertheless this difference in our model, we arbitrarily set [Fgf8a : HSPG$^{\mathrm{cellSurf}}$] to 2× the total ECS Fgf8a concentration. As the spacing between the experimental values does not coincide with the spacing $\Delta y$ between $xz$-planes of the image, we linearly interpolate values between neighboring data points. We convert normalized intensity value to concentrations by multiplying with the maximum concentration of 7.9 nM measured by FCS. The resulting concentration field is shown in Fig. 3b).

### S2 ECS asymmetries are sufficient to explain the dorsal-ventral Fgf8a gradient

In addition to the gradient along the AV axis, Fgf8a forms a weaker gradient along the DV axis within the marginal blastomere with increasing Fgf8a concentration towards the dorsal side, as shown in Fig. S1 at ≈75% epiboly [50]. This gradient could be caused by spatial variations in source or sink rates, or it could be a consequence of DV asymmetry of the ECS geometry with a thicker shield cell layer at the dorsal side. To test whether geometric asymmetry is sufficient to explain the DV gradient, we compute *de novo* 1D concentration profiles along the DV axis for different time points at ≈75% epiboly by averaging concentrations over $yz$-planes in a region 100 μm above the margin (dashed yellow lines in Fig. S2, left).

The resulting concentration profiles (Fig. S2, right) show that a gradient comparable to the experimentally observed one (Fig. S1, right) spontaneously emerges over time and reaches steady state after ≈30 minutes. Since all other model parameters, in particular all sink and source rates, were kept the constant along the DV axis, this shows that asymmetries in the ECS geometry are sufficient to explain the Fgf8a DV gradient. This is in line with the observation that DV concentration profiles are flat in ECS geometries of earlier epiboly stages, where geometric asymmetries are not as pronounced yet (Fig. S3).

While this shows that geometric asymmetries are sufficient, they may not be necessary. Additional factors may also contribute to the DV gradient, such as spatial asymmetries in source rates. For instance, it has been shown in *in vivo* that fgf8a-mRNA concentrations are higher in the cells at the dorsal side [44, 97]. Another factor could be faster growth (and thus advection) rates at the dorsal side, as it has been shown that beyond 50% epiboly, cells at the dorsal side move faster than cells at the ventral side [72].

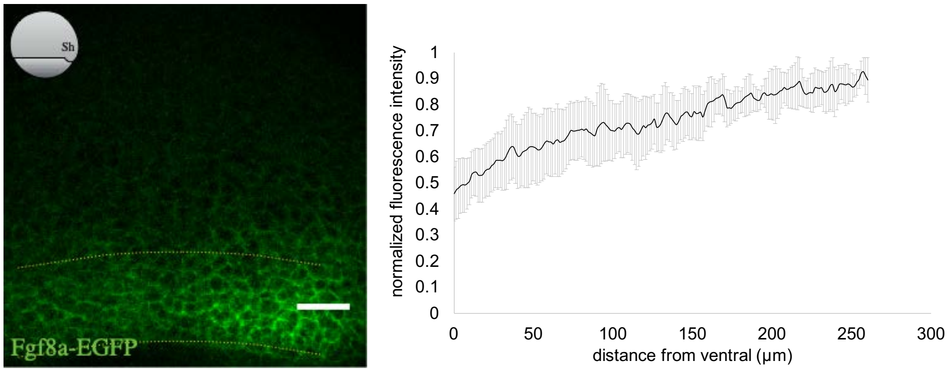


Figure S1: Fgf8-EGFP gradient in the ECS at midgastrula stage (≈75% epiboly). The normalized fluorescence intensity profile in the ventral-to-dorsal direction (right) was extracted from sum-intensity z-projected confocal images (left). Values are intensity averages across a region at the marginal blastomere (dotted yellow lines). Data are represented as mean ± standard deviation. The embryo is oriented as in Fig. 5c) (ventral left, dorsal right, animal pole top, vegetal pole bottom), see inset sketch; scale bar: 50 µm. Figure reproduced with permission from Harish et al. (2023).

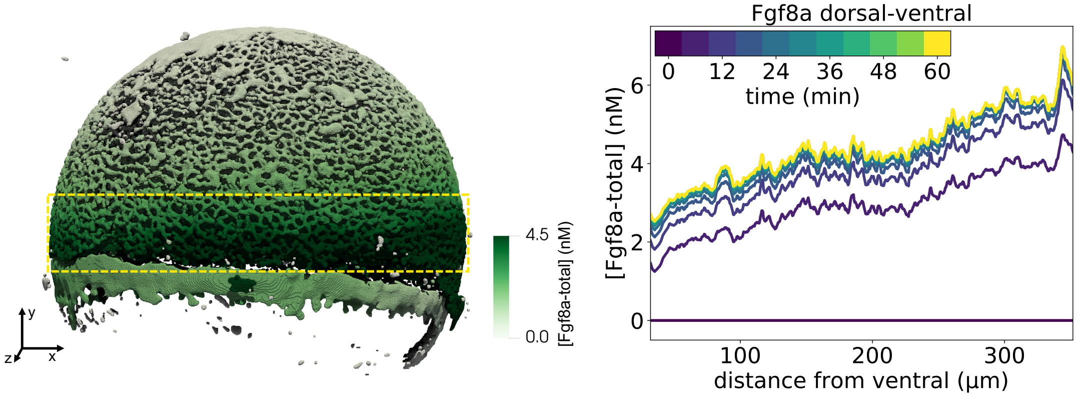


Figure S2: Simulated *de novo* Fgf8a gradient and DV concentration profiles at ≈75% epiboly. The concentration profiles in the ventral-to-dorsal direction (right) were computed by averaging concentration values over $yz$-planes within the marginal region (dashed yellow lines). The model has the same orientation as the embryo in Fig. S1. Color represents concentrations in nM (left) and simulated time of gradient formation (right), see color bars.

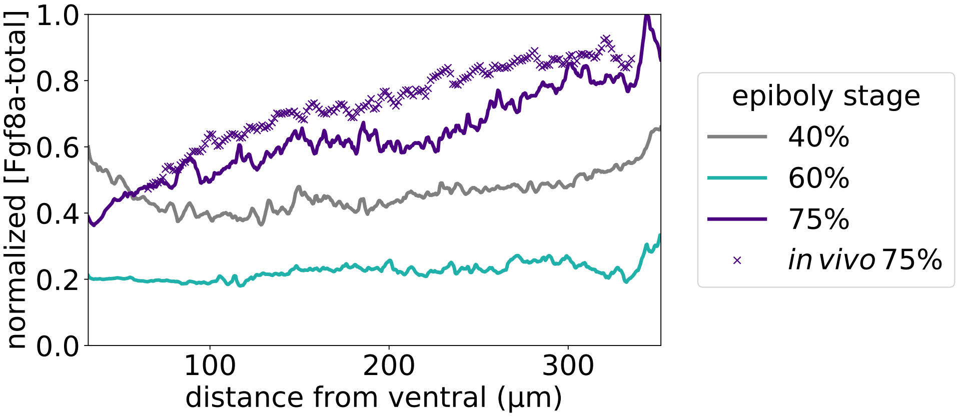


Figure S3: Normalized *de novo* Fgf8a DV concentration profiles at different stages of epiboly (color, inset legend). A gradient similar to the one observed *in vivo* spontaneously emerges at ≈75% epiboly (purple, line: simulation, symbols: experiment). The computationally predicted profiles are flat in geometries of 40% and 60% epiboly (gray and cyan lines, respectively). All profiles are computed at $t = 60\,\text{min}$.

### S3 The Fgf8a normalized gradient is robust to sink function of source cells

In accordance with *in vivo* observations [90, 98], our model assumes that source cells express Fgfrs, i.e., they simultaneously act as sinks. To test whether the sink function of source cells influences the Fgf8a AV gradient, we simulate *de novo* gradient formation as described in Section 3.1 but with [Fgfr] = 0 at all source cells. Figure S4a shows that this increases the absolute peak concentration. This is expected, as less morphogen is degraded in the source region. After normalization, however, the source-is-sink and source-not-sink profiles almost coincide, with a slightly shorter range in the source-not-sink gradient. This shows that the impact of this assumption on the overall simulation results is negligible.

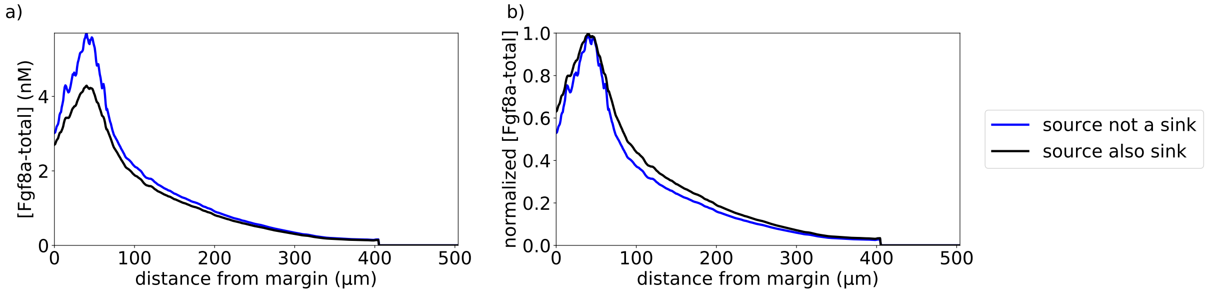


Figure S4: Simulated AV concentration profiles of *de novo* Fgf8a gradient formation when source cells are not sink cells (blue, inset legend) compared to the baseline assumption that sources simultaneously act as sinks (black line). Concentration profiles before (a) and after (b) normalization are shown at $t = 60\,\text{min}$.

### S4 ECS volume is almost constant throughout epiboly

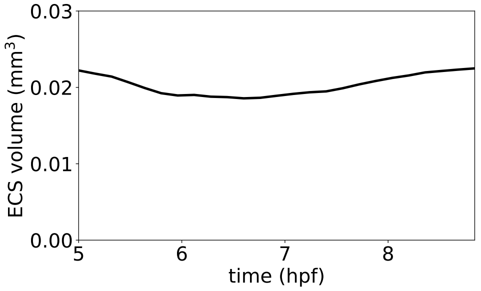


Figure S5: Reconstructed ECS volume at all 25 time point between 5 hpf (≈30% epiboly) and 9 hpf (≈90% epiboly). The volume was computed by summing the volumes of all grid cells inside the ECS.

It has been proposed that tissue growth contributes to gradient formation by changing the diffusion volume, diluting the morphogen concentration [1, 84, 125]. We estimate this dilution factor for Fgf8a gradient formation during zebrafish epiboly by computing the total ECS volume for the 3D geometry reconstructions of all 25 time points of the light-sheet microscopy video ranging from 5 hpf (≈30% epiboly) to 9 hpf (≈90% epiboly). The volume is estimated by summing the volumes all grid cells inside the ECS. This volume likely overestimates the real ECS volume, as it is biased by segmentation errors as discussed in Section 2.5. Nevertheless, as this overestimation error is constant over time, this provides a good estimate of the net change of ECS volume throughout epiboly. We find that the total ECS volume does not increase during the 4 hours of epiboly (Fig. S5). This means that the increase of blastoderm area by spreading over the yolk is counterbalanced by its thinning, thereby maintaining a constant volume. Interestingly, this is although the number of cells increases almost linearly with time between 5 and 9 hpf, approximately doubling from $\approx 10^4$ to $\approx 2 \cdot 10^4$ cells while approximately maintaining the cell density [66]. Increasing the number of cells while maintaining a constant ECS volume suggests that the interstitial space between the cells decreases, and hence, the tortuosity-mediated diffusive hindrance increases throughout epiboly. A constant ECS volume also suggests that the effect of growth-induced morphogen dilution is negligible during epiboly.

### S5 Determining the ECS diffusive tortuosity enables quantifying geometric hindrance

As shown in Section 3.2, the Fgf8a AV gradient is sensitive to changes in the ECS geometry. However, the impact of the geometry may only be indirect through factors such as the total surface areas of source and sink cells, or the total volume (and hence mass) of HSPG. We quantify the diffusive hindrance from the ECS geometry alone by computing the geometric tortuosity of the ECS. Tortuosity $\tau_\mathrm{d}$ is commonly used to quantify geometric hindrance in a porous medium [3, 10, 45, 104, 113]. It is defined as:

$$\tau_\mathrm{d} = \left( \frac{\langle L_\mathrm{d} \rangle}{L_\mathrm{s}} \right)^2 = \frac{D_\mathrm{m}}{D_\mathrm{eff}}\,, \tag{S1}$$

where $D_\mathrm{m}$ is the molecular diffusion coefficient of the morphogen and $D_\mathrm{eff}$ is the effective, coarse-grained diffusion coefficient in the porous medium. This is related to the average path length $\langle L_\mathrm{d} \rangle$ traveled by a diffusing molecule, compared to the straight-line distance $L_\mathrm{s}$.

We determine $\tau_d$ of the 3D ECS geometry at 60% epiboly by simulating a FRAP (fluorescence recovery after photobleaching) experiment. For this, we initialize the lower half of the ECS with a uniform concentration field and set the concentration in the upper half (above 500 µm above the margin) to zero, as shown in Fig. S6a. We then simulate pure diffusion, i.e., without HSPG binding, sources, and sinks, and measure the time required for the concentration field to uniformly distribute throughout the ECS. This time depends on the molecular diffusion constant $D$ and on the modeled ECS geometry. Comparing this time to the one determined in a plain spherical shell (see Fig. S6b) of identical size and with identical $D$, we find $\tau_d = 2.5$.

This value agrees with recent tortuosity estimations of the zebrafish brain ECS at 24 hpf, where the ECS has been reconstructed based on combined fluorescence and transmission electron microscopy images [135]. In this reconstruction, FRAP has been simulated using an agent-based particle Monte-Carlo method, and $\tau_d$ has been estimated to range from 2.0 to 2.7 for different parts of the brain [135].

This means that—excluding other hindrance factors—diffusive transport is less than half as efficient in the ECS geometry than in an empty shell without cells. As shown in Section 3.3, upscaling the ECS tortuosity by decreasing the diffusion coefficient by the factor $\tau_d$ leads to a steeper gradient with a shorter range. This shows that the tortuosity of the ECS geometry is sufficiently high to influence gradient formation by directly hindering morphogen diffusion.

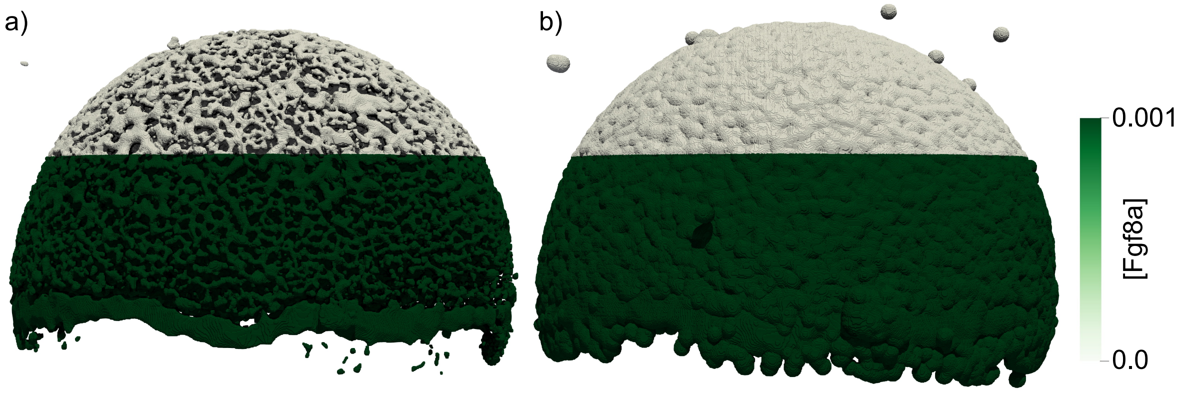


Figure S6: Initial condition of simulated FRAP experiment in the ECS geometry (a) and the shell geometry (b). The lower half of the ECS is uniformly initialized with [Fgf8a] = 0.001 (arbitrary unit), and the upper half (above 500 µm above the margin) with [Fgf8a] = 0. To estimate the diffusive hindrance of ECS tortuosity alone, we simulate diffusion without reactions, measuring the [Fgf8a] total mass recovery in the bleached half. Fitting the recovery curve of the shell geometry to the curve of the ECS geometry, we find the ratio between the local and effective diffusion coefficient, i.e., the tortuosity factor $\tau_d = 2.5$.

#### S6 Supplementary figures

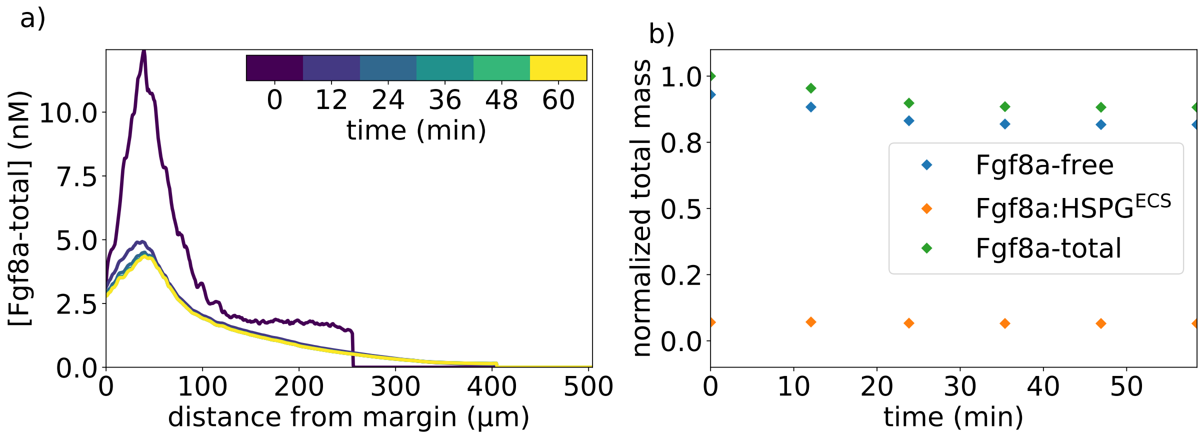


Figure S7: Simulated Fgf8a gradient using the optimized parameters from Tab. 1 with experimental profile as initial condition. a) Absolute AV concentration profiles of Fgf8a-total at different simulated times (color). The experimental profile used as initial condition is shown in black (time 0). b) Normalized total mass of the two Fgf8a fractions Fgf8a : $\mathrm{HSPG}^{\mathrm{ECS}}$ and Fgf8a-free, along with their sum (Fgf8a-total). Although this set of parameters does not maintain the absolute peak concentration, it preserves the *in vivo* Fgf8a / Fgf8a : $\mathrm{HSPG}^{\mathrm{ECS}}$ ratio of 93% / 7%).

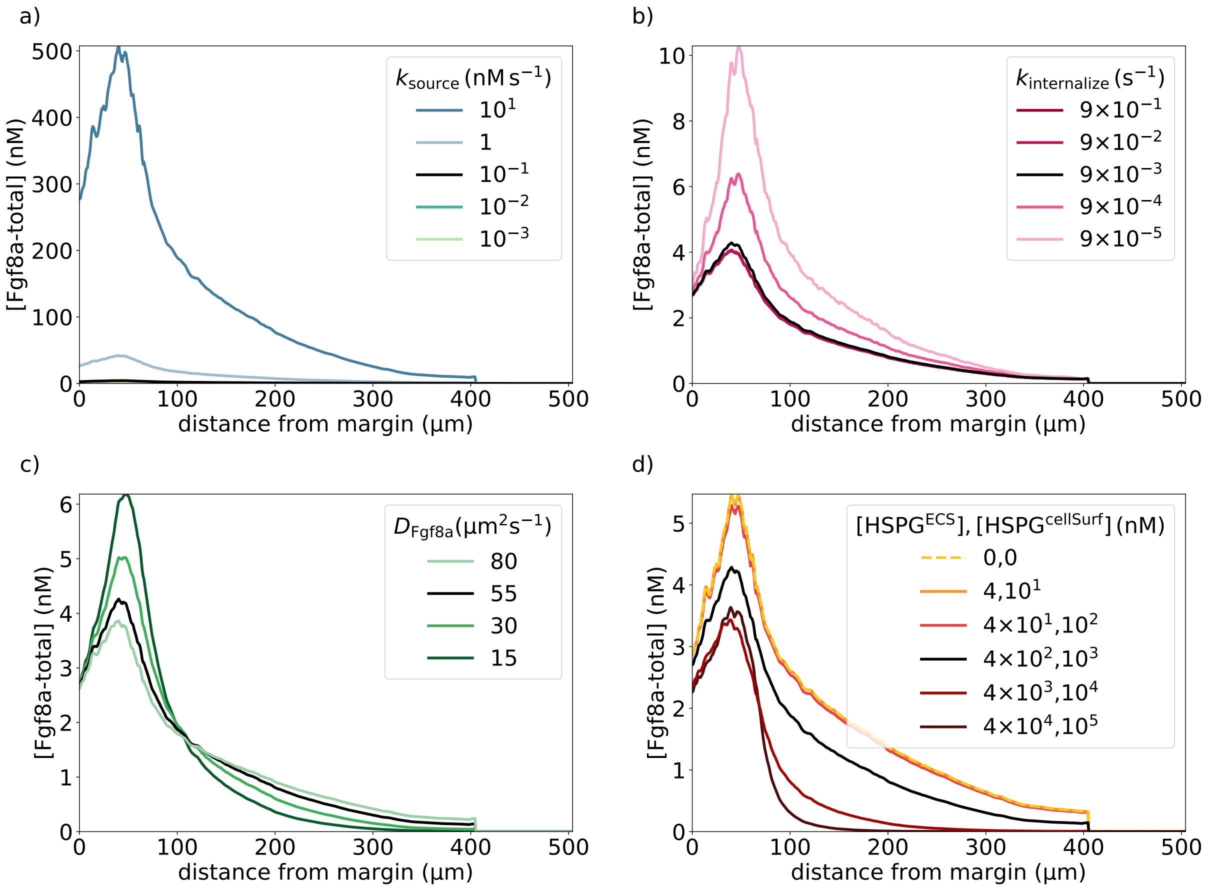


Figure S8: Absolute concentration profiles for the robustness tests of simulated *de novo* Fgf8a gradient formation. Peak concentrations are higher for larger source rates (a) and for lower sink rates (b) (color, inset legend). Gradient profile robustness becomes apparent after normalization with the respective peak concentration (Fig. 7). Changes in the effective Fgf8a diffusivity, by changing either the molecular diffusion coefficient of Fgf8a (c) or the HSPG concentrations (d), strongly affect the gradient shape in addition to the peak concentration. The more HSPG binding and the smaller the diffusion coefficient, the steeper and shorter the gradient. The baseline gradient for the nominal parameters from Tab. 1 is shown as a solid black line in all panels. All profiles are shown at $t = 60\,\text{min}$.

#### S7 ECS reconstructions at all 25 time points

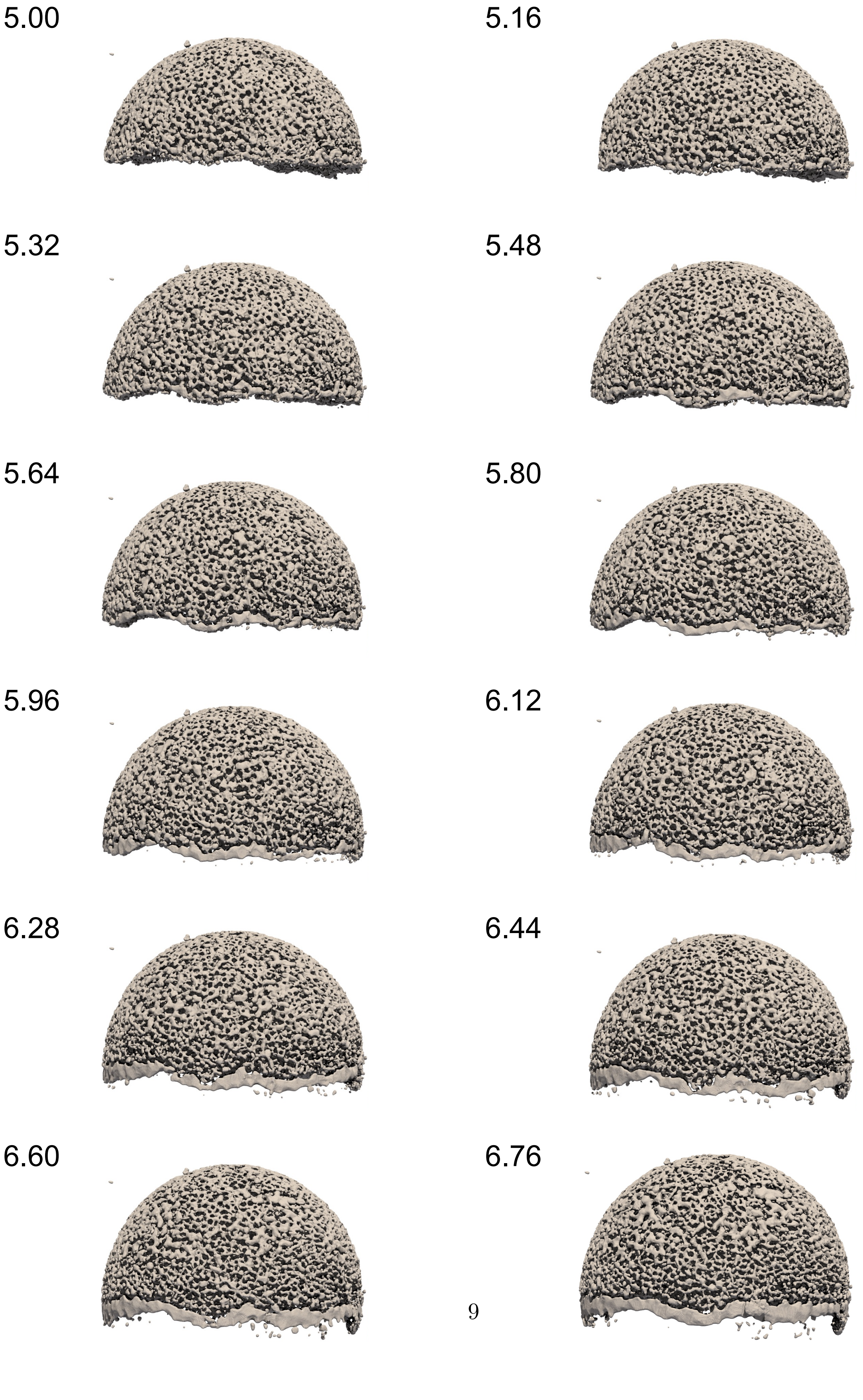
5.00
5.16
5.32
5.48
5.64
5.80
5.96
6.12
6.28
6.44
6.60
6.76

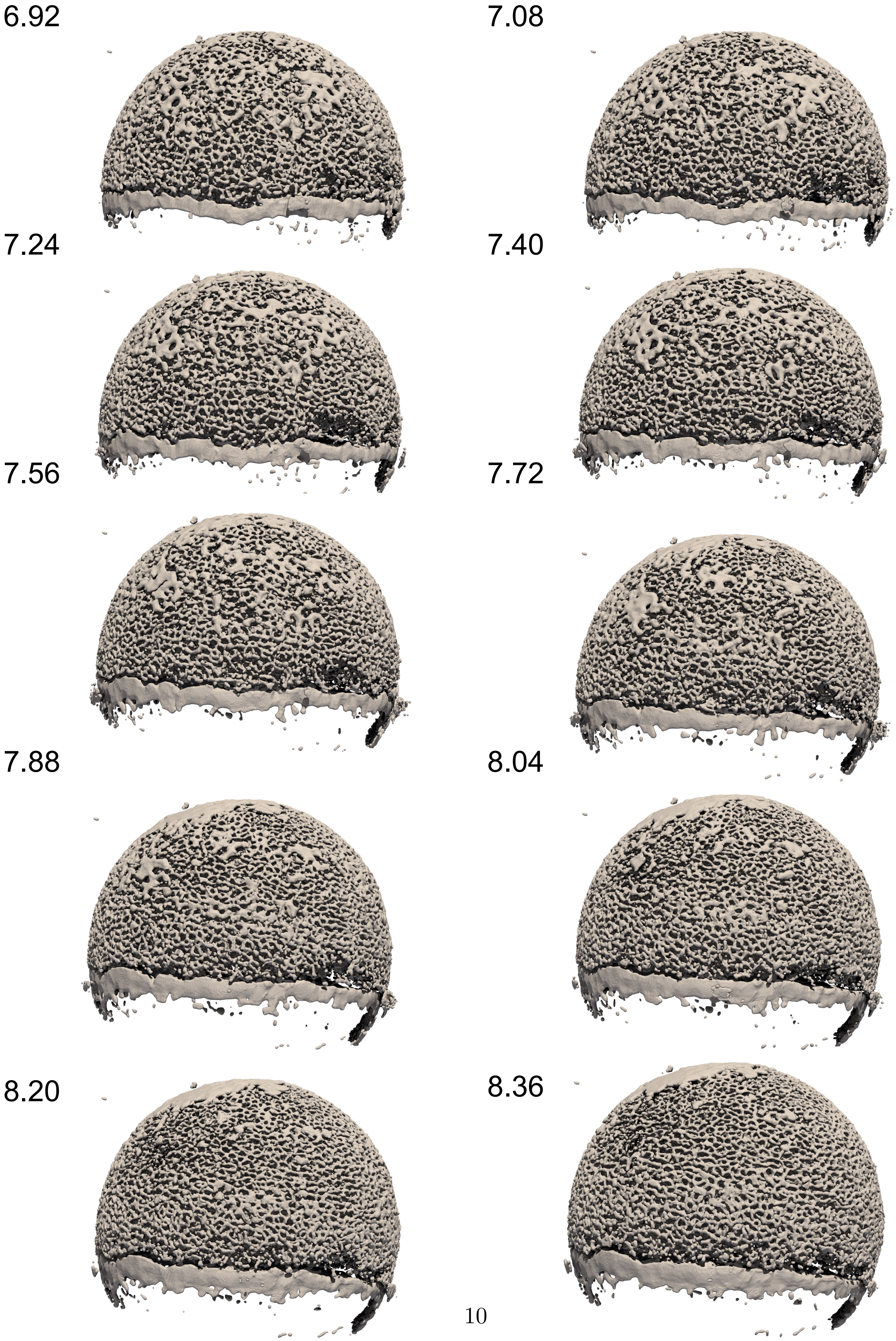
6.92
7.08
7.24
7.40
7.56
7.72
7.88
8.04
8.20
8.36

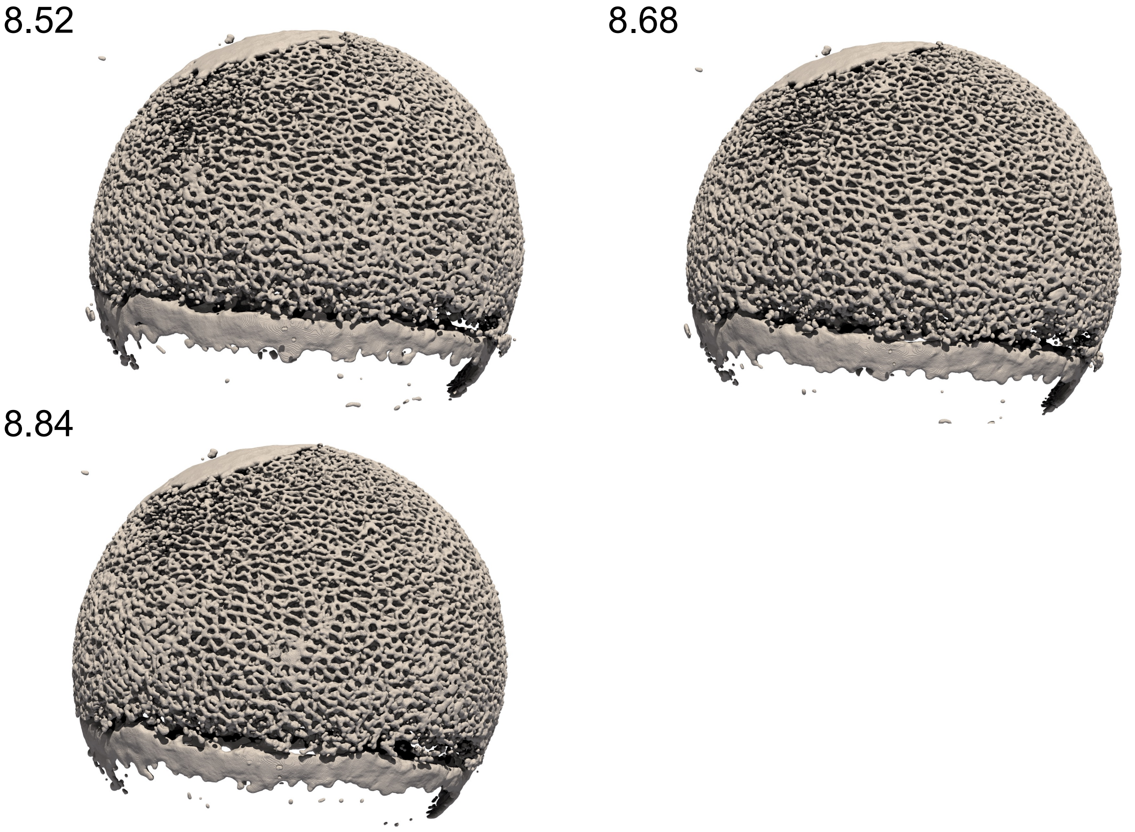


Figure S9: Visualization of the signed-distance-function representation of the ECS boundary at all 25 time points of the light-sheet microscopy video, ranging from 5 hpf (≈30% epiboly) to 9 hpf (≈90% epiboly). For each time point, we show a surface rendering of $\phi_{\mathrm{ECS}} = 0$ in gray after margin alignment. Numbers are the approximate hours post-fertilization at the start of each respective frame acquisition.
